# The abnormal C-terminus in DVL1 impacts Robinow Syndrome phenotypes

**DOI:** 10.64898/2026.02.14.705933

**Authors:** Shruti S. Tophkhane, Gamze Akarsu, Sarah J. Gignac, Katherine Fu, Stephanie Xie, Esther M. Verheyen, Joy M. Richman

## Abstract

Robinow Syndrome is a polygenic, rare skeletal disorder characterized by craniofacial and limb defects. The genes involved are in the Wingless-related Integration site-1 (WNT) pathway and DVL1 (Dishevelled 1) is the most commonly affected gene. In all pathogenic variants of DVL1, a frameshift replaces the C terminus with a novel peptide. We tested whether the variant *DVL1^1519ΔT^* was sufficient to alter development in vivo and in vitro in two animal models. We compared phenotypes to wtDVL1 or DVL1 with a stop codon at position 1519. Misexpression of *DVL1^1519ΔT^* in the developing face of chicken embryos with an avian retrovirus, leads to a widening of the frontonasal mass similar to the human facial phenotype and ultimately to inhibition of skeletogenesis that was also verified in primary cultures of frontonasal mass mesenchyme. In luciferase assays carried out in facial mesenchyme, the wt*DVL1* activated canonical and JNK PCP WNT signalling however the *DVL1^1519*^* and the *DVL1^1519ΔT^* variant removed some but not all of the signaling activity. We also determined that there is mislocalization of the protein expressed from *DVL1^1519ΔT^* in the nucleus while the other two constructs were mainly found in the cytoplasm. In complementary *Drosophila* experiments using a variety of readouts, only the *DVL1^1519ΔT^* variant impacted morphogenesis and signaling. This is the first study to clarify the pathogenesis of Robinow syndrome is due to the novel C-terminus of DVL1 which exerts dominant interference on morphogenesis, skeletogenesis and WNT signaling.

## Introduction

Robinow syndrome is a rare polygenic skeletal disorder associated with variants in the Wingless-related integration site-1 (WNT) signaling pathway (Table S1). Autosomal dominant forms of Robinow syndrome are caused by variants in *WNT5A* (WNT ligand), Frizzled 2 (*FZD2,* receptor) or Dishevelled (*DVL1,* 2, 3, cytoplasmic adaptor protein required for signal transduction) (1). The recessive form of Robinow syndrome is due to homozygous loss-of-function variants in the ROR2 receptor (2, 3). The recessive form of Robinow syndrome is more severe compared to the autosomal dominant forms but the same pathways are involved. DVL is at the crossroads of all of the WNT signaling pathways (4, 5). The canonical WNT pathway begins with WNT ligands binding to co-receptors FZD and LRP. This occupancy of the receptors recruits DVL and AXIN to cell membrane which disrupts the βcatenin degradation complex. Ultimately β-catenin accumulates in the nucleus, thereby triggering gene transcription (6). Non-canonical pathways include the JNK-Planar cell polarity pathway that involves co-receptors FZD and ROR. Once the ligand is bound, there is recruitment of DVL to the cell membrane and signaling via small GTPases leads to activation of ROCK and JNK thereby triggering changes in cell shape, motility and polarity (4). Calcium signaling also involves the FZD receptors and co-receptors (ROR, RYK). Binding of the ligand triggers calcium release and phosphorylation of the NFAT transcription factor that leads to transcriptional changes (4). We have shown that *DVL1* variants affect the balance of canonical and non-canonical signaling (7). Similarly specific variants of *FZD2* affect either canonical or non-canonical WNT signaling (8). We hypothesize that all gene variants that cause Robinow syndrome change WNT signaling in similar ways leading to common phenotypes. The Robinow syndrome primary manifestations are craniofacial defects (jaw hypoplasia, broad nasal bridge and hypertelorism i.e. wide-set eyes, osteosclerosis) and mesomelic limb shortening (9–11). The craniofacial and skull phenotypes predominate in cases of Robinow syndrome caused by DVL variants (1, 12, 13).

*DVL1* is the most commonly affected gene in autosomal dominant-Robinow syndrome (23/67 patients; MIM#616331) (1, 12–15). *Dvl1* is ubiquitously expressed in the mouse embryo (16) and more highly expressed in the eye primordium and in the spinal cord of chicken embryos (17) which does not immediately explain the specific limb and facial anomalies associated with Robinow syndrome.

*DVL1* variants primarily affect the C-terminus of the protein (1, 12, 13, 15, 18). The DVL1 protein signaling functions occur via three highly conserved functional domains - DIX, PDZ, and DEP (Fig 1A) (19). A ‘basic region’ lies between the DIX-PDZ domains and a ‘proline-rich region’ between PDZ-DEP domains followed by the conserved C-terminus (20). The nuclear localization sequence (NLS) and nuclear export sequence (NES) mediate DVL shuttling which is critical for WNT signaling (21, 22). The functional domains and the C-terminus have been shown to regulate WNT signal transduction and ubiquitination (19–27). The terminal 23 amino acids of C-terminus have an important function in the regulation of balance between the canonical and non-canonical signaling (20).

**Figure 1.**
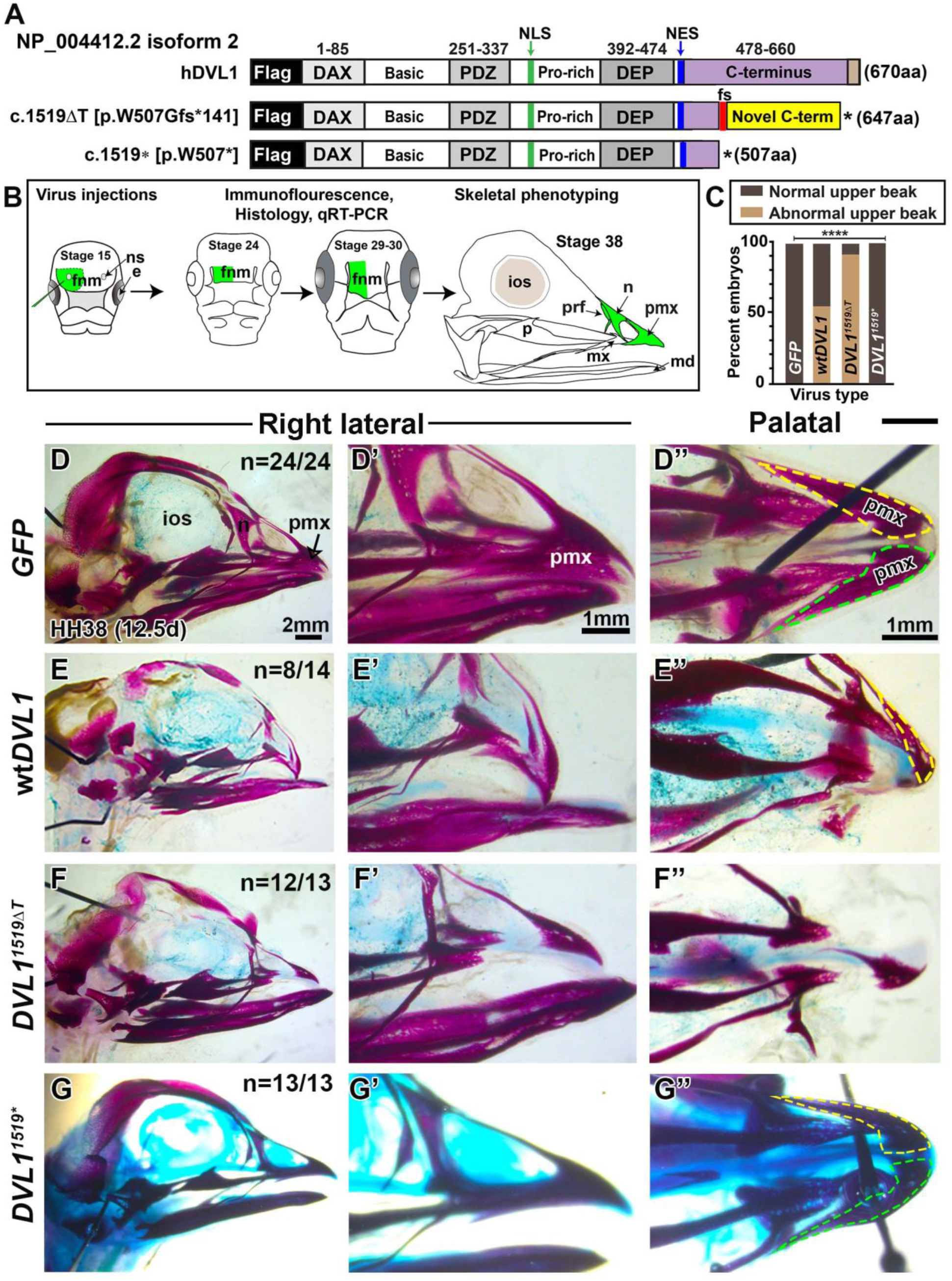
Experimental design and effects of DVL1 variants on skull development. A) Wild-type human DVL1 with an N-terminal flag tag, followed by the main signaling domains followed by the C-terminal. The DVL1^507fs*141^ (DVL1^1519ΔT^) and truncating construct DVL1^507*^(DVL1^1519*^) also have an N-terminal FLAG-tag. The nuclear localization sequence (NLS, IxLT) and nuclear export sequence (NES, M/LxxLxL) are maintained in all the constructs. However, the DVL1^1519ΔT^ construct has a frameshift (fs, red), leading to a novel C-terminus of 141 aa, followed by a STOP codon (asterisk). B) Schematic of RCAS virus injection and skeletal phenotyping. Embryos were injected in the right frontonasal mass at HH15 (embryonic day 2.5) and phenotyping was performed at multiple stages. C) Contingency analysis followed by Wilson/Brown (fraction of total) test showing a statistically significant difference in the fraction of total embryos with abnormal upper beak. D-D’’) Wholemount skulls stained with alcian blue and alizarin red showed a normal patterning of frontonasal mass derived bones (premaxilla, nasal, prefrontal) in GFP-injected specimens (24/24). E-E’’) In contrast, embryos injected with wild-type DVL1 (n=8/14) had a shorter, deviated beak with missing premaxilla. F-F’’) Variant hDVL1 (n=12/13) displayed hypoplastic premaxillary bones. G-G’’) Normal beak formation in all embryos injected with the 1519* truncated variant.. Key: e- eye, fnm – frontonasal mass, fs – frameshift, ios – intraorbital septum, md – mandibular bone, mx- maxillary bone, n- nasal bone, NES – nuclear export sequence, NLS – nuclear localization sequence, ns – nasal slit, p – palatine bone, pmx – premaxilla, Scale bars in D-D’’) apply to all images.

Currently there are 15 *DVL1* coding variants that are pathogenic (ClinVar). These variants result in a frameshift that replaces most of the C-terminus with a novel sequence (Table S2) (1, 12, 13, 15). The frameshifts switch to the same alternative reading frame. The frameshift that occurs at c.1519delT results in a total of 141 abnormal amino acids (Fig. 1A). There is no effect on the sequence of the DIX, PDZ, DEP domains or the NLS. The looping of the C-terminus to the PDZ domain (20) is unlikely to occur due to replacement of the terminal 23 amino acids. The role of the novel C-terminal peptide on WNT signal transduction and protein stability is unclear. We have shown that the autosomal dominant Robinow Syndrome-*DVL1^1519ΔT^* variant dominantly supresses the activity of wt*DVL1* resulting in lower canonical WNT signaling (7). In addition, *DVL1^1519ΔT^* protein showed significantly increased nuclear localization suggesting that the NES (follows the DEP) domain may be affected (7).

One copy of the variant gene is sufficient to cause autosomal dominant Robinow syndrome. Heterozygous mice with loss-of-function mutations in *Wnt5a* (28), *Dvl1, Dvl2, Dvl3* (29), and *Fzd2* (30) are phenotypically normal and therefore do not phenocopy Robinow syndrome. *Dvl1^-/-^*mice are born and survive until adulthood without skeletal abnormalities (31). *Dvl1^-/-^*, *Dvl2^-/-^* mice have severe skeletal malformations and neural tube closure defects (32), while *Dvl1^-/-^*, *Dvl3^-/-^*mice die of unknown causes (33). Loss-of-function experiments are insufficient to understand the pathogenesis of autosomal dominant Robinow syndrome. Instead, the variants need to be tested to see whether they are sufficient to alter development. Our group has shown that the misexpression of autosomal dominant Robinow Syndrome variants in WNT5A produces jaw shortening (34) and shorter limb skeletal elements (35) due to disruption of cell polarity and less secretion of the protein. *FZD2* variants are sufficient to inhibit ossification of bones and interfere with cell signaling (8). *DVL1* variants are sufficient to cause limb cartilage to be disorganized, to disrupt wing patterning in flies and to change the balance of canonical WNT and JNK-PCP signaling (7, 36). Here we are studying the midface which is affected in nearly all individuals with Robinow syndrome, irrespective of genotype. We examined the effects of *DVL1^1519ΔT^* on upper beak (chicken) morphogenesis, cellular dynamics, chondrogenesis, gene expression, WNT signal transduction, and protein localization.

To characterize the specific functions due to the novel C-terminal peptide, we introduced a stop codon (TAA) after nucleotide *1519* (c.*DVL1^1519*^*codes for p.DVL1^507*^)(12). We introduced this truncated version of the *DVL1* gene into chicken and *Drosophila*. Interestingly, removal of the novel C-terminus allowed normal beak morphogenesis indistinguishable from expression of *GFP* control. This was supported by absence of wing and body defects in *Drosophila* by misexpression of *DVL1^1519*^*. Our in vivo data in both animals showed that *DVL1^1519ΔT^* disrupts cell fate and significantly inhibits chondrogenesis at early stages of development compared to wt*DVL1* and *DVL1^1519*^*. Overall, this study demonstrates that the novel C-terminal sequence in *DVL1^1519ΔT^* dominantly interferes with protein function leading to the craniofacial and cell signaling defects. The abnormal C-terminus removes some signaling function and interferes with other aspects of development. Together these changes in function due to the abnormal C-terminal peptide contribute to the facial pathogenesis of Robinow syndrome.

## Results

In order to determine whether there were specific effects due to the abnormal C terminus of *DVL1^1519ΔT^* we misexpressed human *DVL1^1519ΔT^* in the chicken face or in *Drosophila* under the control of specific GAL4 drivers (7, 36). The injections of the avian virus were carried out after neural crest cells have migrated into the face (stage 15; Fig. 1B). Results were compared to the effects to wild-type (wt) *DVL1*, or *DVL1^1519*^* where the C terminus past the frame shift is not translated into protein (Fig. 1A)(12). The negative control for chicken viruses was GFP virus which does not affect development (34, 35, 37). We confirmed that the avian retroviral titre was similar in all three h*DVL1*-containing viruses (Fig. S1A-D). In addition, the three forms of DVL1 were translated into proteins of the expected sizes in chicken cells (Fig. S2A,B) (13) and in *Drosophila* cells (Fig. 2F).

**Figure 2.**
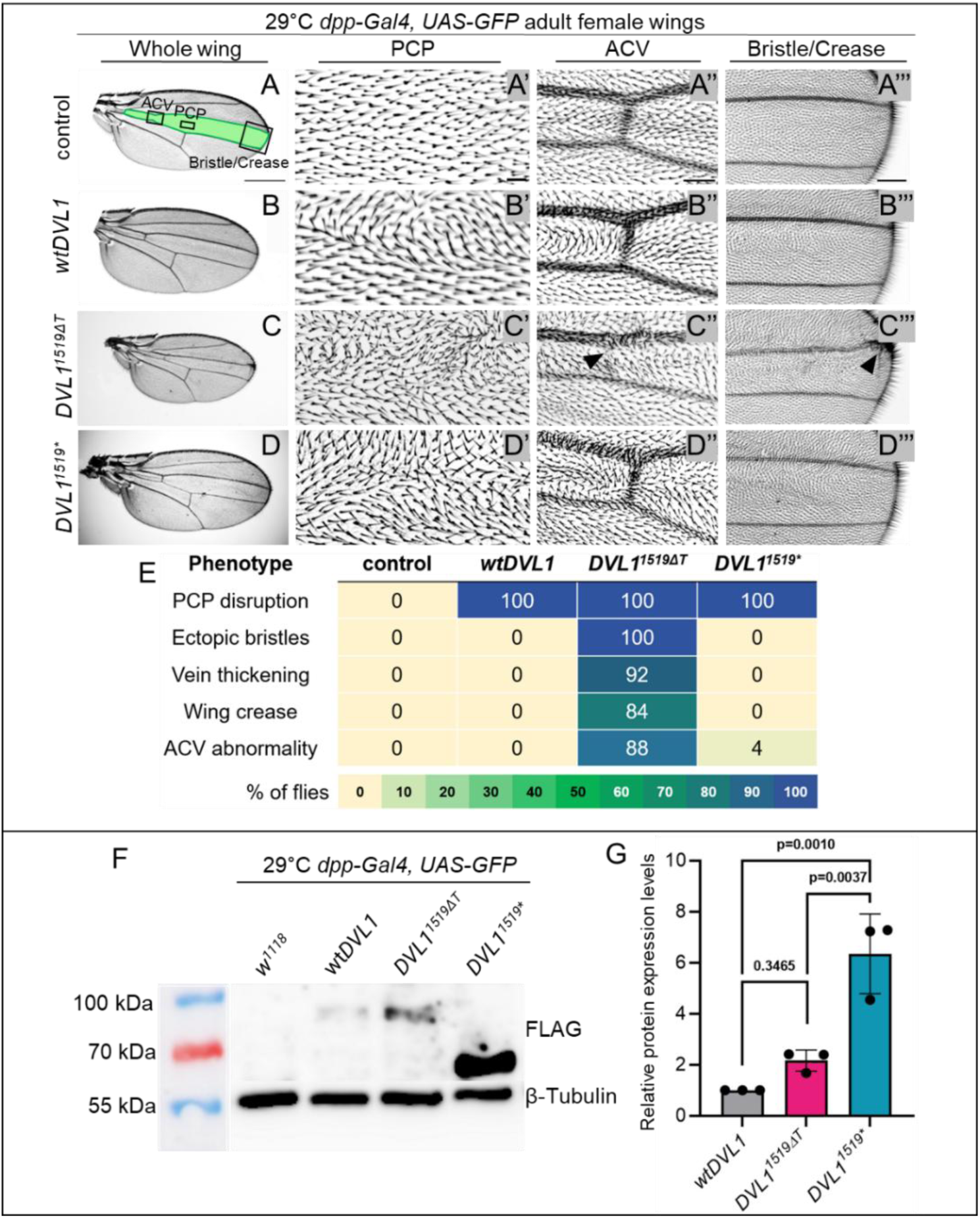
Adult wing phenotypes induced by the expression of DVL1 transgenes in the *dpp-Gal4* domain and protein expression levels of L3 larval heads at 29°C. (**A**) Control wild-type wing with shaded *dpp*-*Gal4* expression domain (green) and black boxes that correspond to zoomed-in views presented in panels (**A’-D’’’**). (**B-D**) Representative *dpp>DVL1-*expressing adult female wings at 29°C. (**A’-D’**) Zoomed in views of PCP defects within a fixed region above the posterior cross vein for control (**A’**) and *DVL1*-expressing (**B’-D’**) adult female wings (**A’’-D’’**). Zoomed in views of anterior cross vein (ACV) in control (**A’’**) and *DVL1*-expressing (**B’’-D’’**) adult female wings. Arrowhead in **C’’** points to reduction of ACV. (**A’’-D’’**) Zoomed in view of adult wing between the L3 and L4 veins in control (**A’’’**) and *DVL1*-expressing **(B’’’-D’’’**) female wings from 29°C crosses where extra bristles and creases are indicated with arrowheads in **C’’’**. Phenotypic frequencies are quantified in (**E**). 25 wings were scored per genotype across n=2 independent experiments. Scale bar = 500 μm (**A-D**), 20 μm (**A’-D’**), 50 μm (**A’’-D’’**), 100 μm (**A’’’-D’’’**). (**F**) Western blot analysis of FLAG tagged wildtype, variant and C-terminal truncated DVL1 protein levels from larval head protein extracts. β-tubulin was used as a loading control. (**G**) On the right, plot of DVL1 protein levels in variant DVL1-expressing tissue relative to control (*dpp*>wt*DVL1*) tissue. Each dot on the bars represents one blot. Error bars show mean with SD. Statistics were performed with a one-way ANOVA test.

The chicken viral and *Drosophila* genetic effects are interpreted as follows: (1) If there are phenotypes caused by wt*DVL1* but no phenotypes caused by misexpression of the *DVL1* variant this indicates a loss-of-function, (2) if the phenotypes are similar to the wt*DVL1*, then the variant has similar functional impacts, (3) if the phenotypes are similar but more severe compared to wt*DVL1*, then the variant causes a gain-of-function, and (4) if the variant phenotypes are neomorphic, the variant has dominant interference effects. The effects of *DVL1^1519*^* were analyzed in similar manner. Through in vivo and in vitro comparisons, we tested two possibilities – that the presence of the novel C-terminal peptide is driving the phenotypes or that the absence of the endogenous C-terminal contributes to Robinow syndrome phenotypes.

### The novel C-terminus in Robinow syndrome variant *DVL1^1519^***^Δ^***^T^*causes upper beak defects via gain-of-function mechanism

In Robinow syndrome, the presence of hypertelorism and a broad nasal bridge often suggests an issue with the development of the medial nasal prominence. The mammalian medial nasal prominences contribute to midline facial structures encompassing nasal septum, nasal bridge, philtrum and midline of the upper lip including the premaxilla (38). We targeted the *DVL1* viruses to the early frontonasal mass at stage 15 that will give rise to the premaxilla and prenasal cartilage (Fig.1B). To directly access the effects of *DVL1* viruses, we collected embryos at stages 24 [48h post injection, E4.5], stage 29-30 [96h post injection, E6.5] and stage 38 (E12.5), covering the period from early morphogenesis (for molecular analysis) to skeletal maturation (Fig. 1B, Table S3).

As expected, injection of *GFP* encoding virus allowed normal upper beak development supported by a normal skeleton (Fig. 1C, D-D’’). Embryos injected with wt*DVL1* and *DVL1^1519ΔT^* viruses displayed a shortened and deviated upper beak (Fig. 1C,E-E’’, F-F’’; Fig. S3B-G). The penetrance of premaxillary phenotype was 57% in wt*DVL1* and 92% in *DVL1^1519ΔT^* (Table S4). We did not observe qualitative differences in alizarin red staining suggesting that the viruses did not affect the ossification of the upper beak bones. In contrast, 100% of embryos injected with *DVL1^1519*^*were normal (Fig. 1G-G’’, Fig. S3H-J). One possible explanation is that the protein translated by *DVL1^1519*^*is non-functional. In *DVL1^1519ΔT^*, the C-terminus is substituted with a unique extended amino acid sequence, leading to the hypothesis that the presence of this peptide causes abnormal upper beak development and suggests that the novel C-terminus causes a gain-of-function compared to *DVL1^1519*^*.

### Similar Planar Cell Polarity phenotypes caused by *wtDVL1, DVL1^1519ΔT^* and *DVL1^1519*^* in *Drosophila* wings

In adult *Drosophila* wings the wing blade is covered with actin hairs called trichomes, which are parallel to each other and pointing distally. This orientation is governed by the Wnt PCP pathway (39, 40). Wing hair polarity defects have been a well-established method for identifying disruptions in the larval and pupal PCP pathway. In our earlier studies, it was shown that all transgenes encoding wildtype and Robinow syndrome-associated DVL1 variants (including the 1519ΔT variant) induced defects in planar cell polarity in adult wing blades (7). Our results showed that the C-terminally truncated DVL1 caused similar PCP defects to the other human DVL1 constructs, in all the wings scored (Fig. 2A-E, n=25 for all). Therefore, the protein coded by *DVL1^1519*^* retains activity in *Drosophila*.

We previously described neomorphic phenotypes only in the wings of the flies that express the *DVL1* variants *dpp>DVL1^1519ΔT^*, *dpp>DVL1^1529ΔG^ dpp>DVL1^1615ΔA^* (7). We found neomorphic phenotypes for the *1519ΔT* variant including thickening in veins, ectopic bristle formation on the edge of the L3 vein, alterations or absence of anterior cross vein (ACV) and a wing crease between the intervein region between L3 and L4 longitudinal veins (Fig. 2E)(7). These mutant phenotypes were not observed in *dpp>wtDVL1* wings or *dpp>DVL1^1519*^*, even with elevated protein levels at 29°C (Fig. 2B’’,B’’’, D’’,D’’’,E). We confirmed that all DVL1 transgenes were expressed in our experimental conditions and found abundant protein, especially for 1519* (Fig. 2F, G). Thus, we conclude that these phenotypes were novel effects of the 1519ΔT variant (7). Unlike the chicken beak experiments, there is activity in the *dpp>DVL1^1519*^* transgene based on the effects on *Drosophila* bristle polarity.

### Lack of C-terminus does not induce abnormal morphology in adult fly tissues

Since the region corresponding to the *dpp-Gal4* expression domain in the adult wing is narrow, we used *hh-Gal4* that span a larger region of developing discs to look for adult phenotypes (36). Flies with the wtDVL1 construct expressed in the hh domain eclose normally had have similar morphogenesis compared to controls (Fig. 3A-B’). Flies that express the *DVL1^1519ΔT^* variant in the *hh* domain at 29°C had difficulties eclosing from their pupal cases due to their abnormal leg morphology, as most of them died trying to eclose or they fell on the food after eclosion (36)(Fig. 3C,C’). We observed that the C-terminal truncated *DVL1^1519*^* expressing flies developed similarly to the *wtDVL1* expressing flies with no dramatic alteration in their wing or leg morphology (Fig. 3D,D’). This experiment again showed that the altered morphology caused only by the expression of the *DVL1* variant, and it is due to the presence of the novel peptide sequence at its C-terminus, not the lack of the highly conserved C-terminus. These fly phenotypes are similar to the chicken beak experiments where loss of the C-terminus had no effect on morphogenesis. However, before we can conclude there is no residual activity the truncated form of *DVL1^1519*^* in the chicken model, more experiments are needed in a variety of contexts.

**Figure 3.**
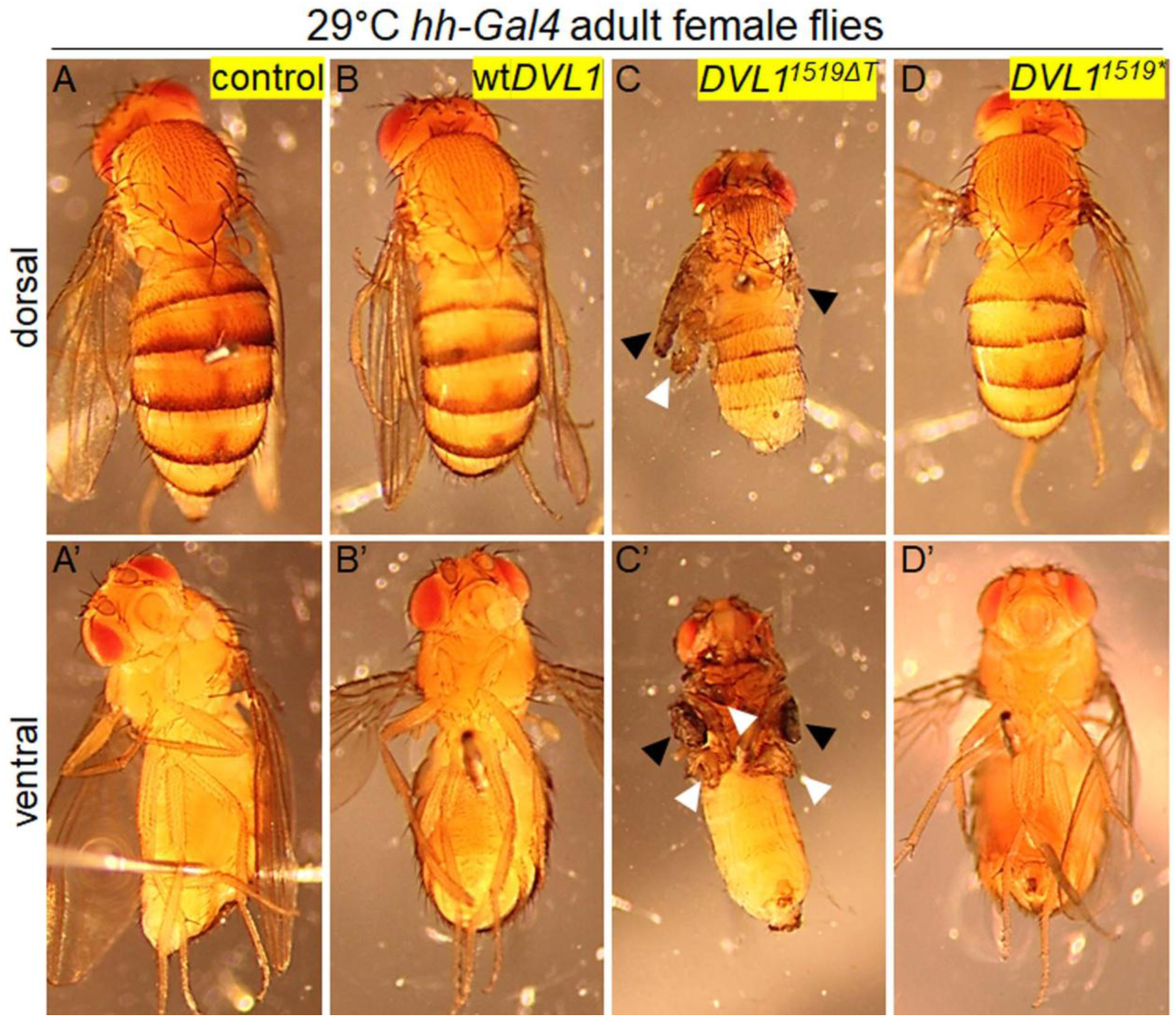
Expression of C-terminally truncated DVL1 does not induce abnormal adult structures when expressed in the *hh-Gal4* domain and summary of signaling defects caused by the wtDVL1, the *DVL1^1519ΔT^* and *DVL1^1519*^* constructs in chicken and *Drosophila*. (**A-A’**) 29°C control adult female fly phenotypes. (**B-D’**) Representative dorsal (A-D) and ventral (A’-D’) view images of female adult phenotypes from *hh>*Gal4 crosses performed at 29°C crossed to (B-B’) wt*DVL1,* (**C-C’**) *DVL1^1519^*^Δ^*^T^*, and (**D-D’**) *DVL1^1519*^*. Arrowheads in **C**-**C’** points to malformed wings (black) and the legs (white) in *hh>DVL1^1519^*^Δ^*^T^* flies. 30 flies were collected and observed from n=4 genetic crosses.

### Robinow syndrome variant *DVL1^1519^***^Δ^***^T^* disrupts differentiation of the prenasal cartilage

To identify the origins of the beak phenotype, we collected embryos at stage 24-25 (E4.5) and stage 29-30 (E6.5-7) for histology, immunostaining and qRT-PCR (Fig. 4A-P; Fig. 5A-R; Tables S6, S7). All the specimens were tested for viral spread using GAG antibody and only those embryos with adequate virus were included in the study (Fig. 4D-F; Fig. 5D-F). We confirmed that all viruses were expressed at similar levels with qRT-PCR (Fig. 4N). Embryos injected with *GFP* and wt*DVL1* had normal frontonasal mass morphology at stage 24 (Fig. 4A-L). There were no differences in SOX9 expression, a marker of prechondrogenic mesenchyme (Fig. 4G-I) and no differences in proliferation (Fig. 4J-M). There was no evidence of a feedback loop whereby gallus DVL1 was affected (Fig. 4O). Genes in the WNT pathway and skeletogenic genes were also not affected (Fig. 4P).

**Figure 4.**
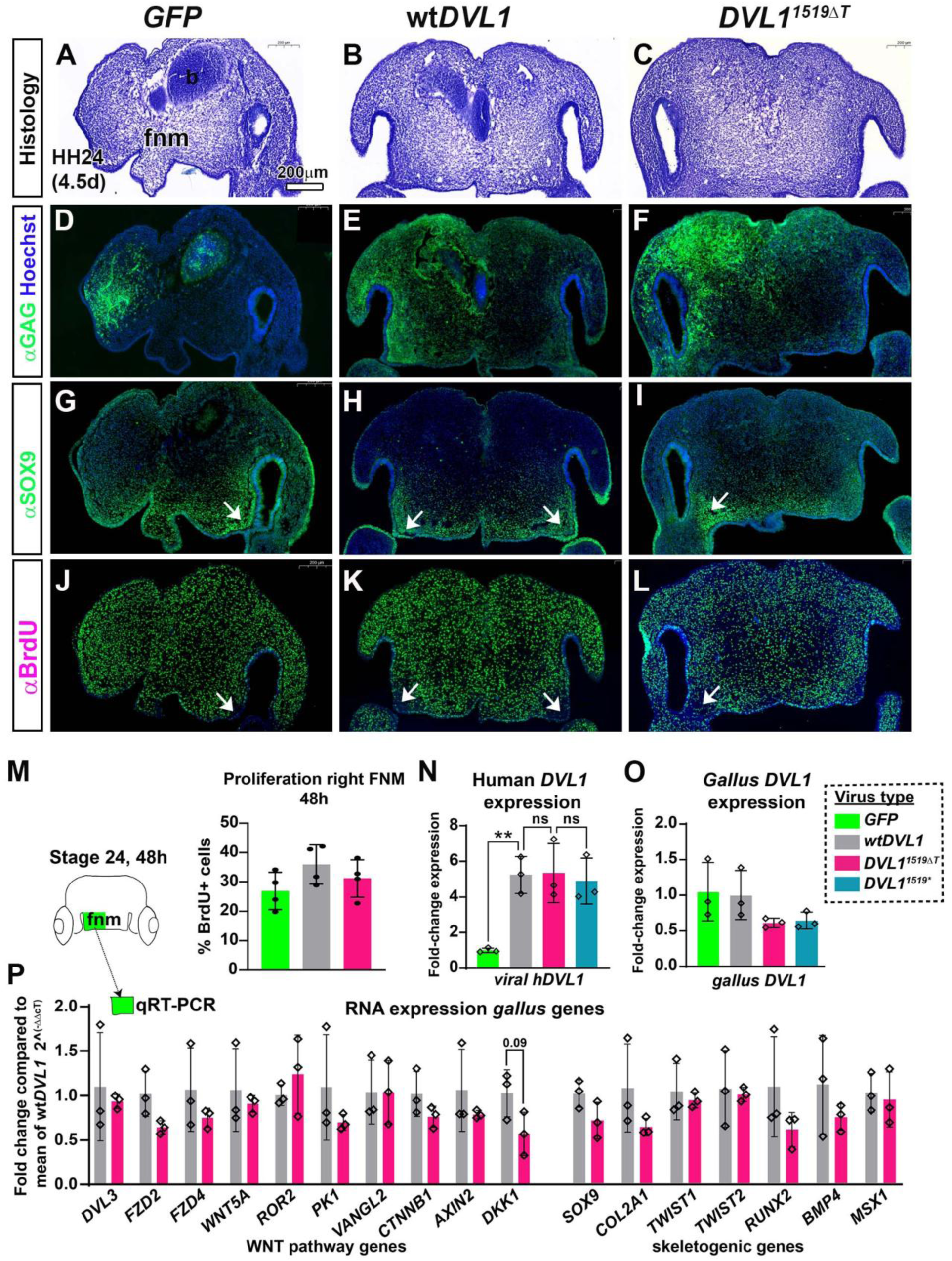
Effects of DVL1 viruses on frontonasal morphogenesis 48h post infection. A-C) Histology of embryos that were used for subsequent immunofluorescence experiments in D-L. D-F) Presence of the viral GAG protein is shown with anti-GAG staining in the frontonasal mass There is more staining on the right side of the frontonasal mass since embryos lie with their right side facing upwards in the egg and this is the most accessible site for injection. G-I) Embryos with staining for SOX9 in the mesenchyme. Some areas of relatively higher expression are seen in the globular processes (arrows). J-L) Proliferation in the mesenchyme was high and not affected by the presence of the DVL1 viruses (compared to areas of GAG staining shown in D-F). M) quantification of proliferation in the right half of the frontonasal mass (see schematic) shows no difference between any of the viruses. N) The levels of human DVL1 expression are about 4-5 fold higher than GFP infected controls. There is no significant difference between the wt,DVL1, 1519ΔT and 1519* viruses. O) There is no change in expression of the gallus *DVL1* gene so no feedback loops are activated. P) Neither genes in the WNT pathway nor those involved in skeletogenesis were significantly altered in the 1519ΔT variant relative to human wt*DVL1* infected frontonasal mass mesenchyme.

**Figure 5.**
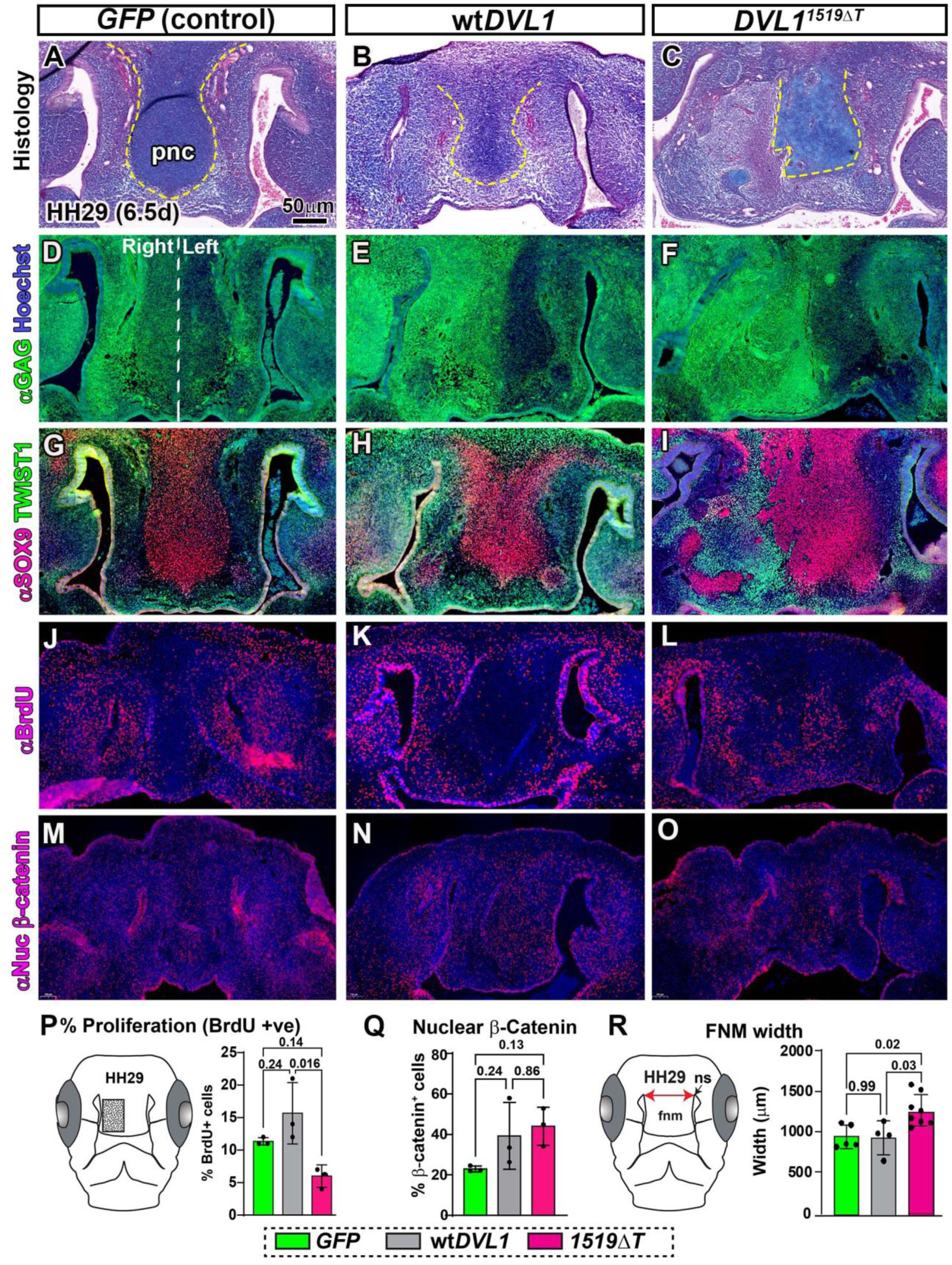
Analysis of embryos injected with GFP, wt hDVL1 and hDVL1^1519ΔT^ at HH29 (embryonic day 6.5): Embryos were injected at stage 15 and fixed 96h post-injection when they had reached stage 29. Near-adjacent sections were used for A,D,G,J,M; B,E,H,K,N; C,F,I,O. Panel L is a different embryo than the others in the right column. A,B) sections stained with Alcian blue and Picrosirius red show differentiating prenasal cartilage in the midline. C) the DVL1 variant has disrupted the cartilage and replaced it with mesenchymal cells. D-F) Broad spread of the virus as shown by GAG staining. G-I) Anti-SOX9 (stains chondrocyte nuclei (red)) and anti-TWIST1 (stains undifferentiated mesenchymal cell nuclei, green). (G, H) In GFP and wild-type hDVL1 infected embryos, SOX9 is expressed in the prenasal cartilage surrounded by undifferentiated TWIST1-positive mesenchyme. I) In embryos infected with hDVL1^1519ΔT^ variant, TWIST1 replaced SOX9 expression in the prenasal cartilage (right side). J-L*, P*) Embryos with BrdU stainin*g* showed significantly lower proliferation in the hDVL1^1519ΔT^ infected area compared to the wtDVL1-infected embryos (P). M-O*,Q*) Nuclear β-catenin is expressed in the prenasal cartilage with no significant differences in expression (Q). R) width between the nasal slits is significantly greater in the variant DVL1 infected frontonasal mass compared to GFP or wtDVL1 infected embryos. Key: fnm – frontonasal mass, ns – nasal slit, pnc – prenasal cartilage. Scale bar = 50µm and applies to all panels.

While *DVL1^1519ΔT^* embryos were initially comparable to wt*DVL1* at stage 24, by stage 29, there was inhibition of the prenasal cartilage (Fig. 5A-C). To determine if the *DVL1^1519ΔT^* virus affected cell fate specification at stage 29, we co-stained sections with SOX9 and TWIST antibodies. In *GFP* or wt*DVL1* injected embryos SOX9 marked the prenasal cartilage while antibodies to TWIST (recognizes TWIST1 and 2) marked the surrounding undifferentiated mesenchymal cells (Fig. 5G,H). In *DVL1^1519ΔT^* embryos, a portion of SOX9-positive prenasal cartilage was replaced by undifferentiated TWIST-positive mesenchyme (Fig. 5I). Notably, the TWIST-positive mesenchyme colocalized with *DVL1^1519ΔT^* virus (Fig. 5I, F). These results suggest the *DVL1^1519ΔT^* virus prevents neural crest-derived mesenchymal differentiation into cartilage that forms the template of the upper beak. At stage 29, the *DVL1^1519ΔT^* virus injected samples showed significantly inhibited cell proliferation compared to wt*DVL1* (Fig. 5J-L,P). Areas where chondrogenesis was present had lower proliferation (compare Fig. G-I to Fig. 5J-L). Since *DVL1* is a major player in the WNT pathway, we examined expression of nuclear β-catenin (marks active canonical WNT signaling). We did not observe any difference in the number of β-catenin-positive cells between the viruses suggesting that the chondrogenic inhibition does not require a change in the levels of canonical WNT signaling (Fig. 5M-O, Q).

Between stages 24-29, the frontonasal mass undergoes narrowing pertaining to the convergence in the mediolateral and extension in the craniocaudal directions (41). Since individuals with Robinow syndrome have a “wide face” phenotype, we measured the width of the frontonasal mass (distance between nasal slits, illustrated in Fig. 5R). Interestingly, frontonasal mass width was significantly greater in the *DVL1^1519ΔT^* embryos compared to *GFP* and wt*DVL1* (Fig. 5R), implying that the *DVL1^1519ΔT^* virus interferes with convergence-extension movements during face development.

We also tested whether the truncated version of DVL1 coded by *DVL1^1519*^* was sufficient to change beak morphogenesis at stage 29. Here the embryos developed normally including forming the prenasal cartilage (Fig. S4A), normal proliferation (Fig. S4C) and no increase in apoptosis (Fig. S4D). These results are consistent with the beak phenotypes and suggest a lack of function for the *DVL1^1519*^* gene.

### *DVL1^1519^***^Δ^***^T^* inhibits chondrogenesis in frontonasal mass micromass cultures

To further explore the direct effects of the 3 forms of DVL1 (wild-type, variant and truncated) on the steps of chondrogenesis, we infected primary cultures of frontonasal mass mesenchyme with the viruses and grew the cells at high density (micromass cultures). On days 4-6 of culture, there were no differences in area occupied by Alcian blue-positive frontonasal mass cartilage sheets (Fig. S5A-H). Thus, the initial condensation of cartilage proceeds normally. Growing the cultures until day 8 revealed that *DVL1^1519ΔT^* virus significantly inhibited expansion of chondrogenesis compared to *GFP,* wt*DVL1* and *DVL1^1519*^*(Fig. 6E-H, C). To delve deeper into the impact of the viruses, a subset of day 8 cultures was sectioned transversely (Fig. 6B). The measurements conducted on the thickest areas within the cultures further clarified effects of the variant *DVL1* on chondrogenesis. Cultures infected with *DVL1^1519ΔT^* were significantly thinner compared to *GFP* and wt*DVL1* (Fig. 6I-L, C). On the other hand, *DVL1^1519*^*infected cultures had thickness comparable to *GFP* and wt*DVL1* (Fig. 6I-L, C). We also confirmed that the differences in the cartilage area and culture thickness could not be attributed to lower expression of human or gallus *DVL1* transcript (Fig. 6M,N).

**Figure 6.**
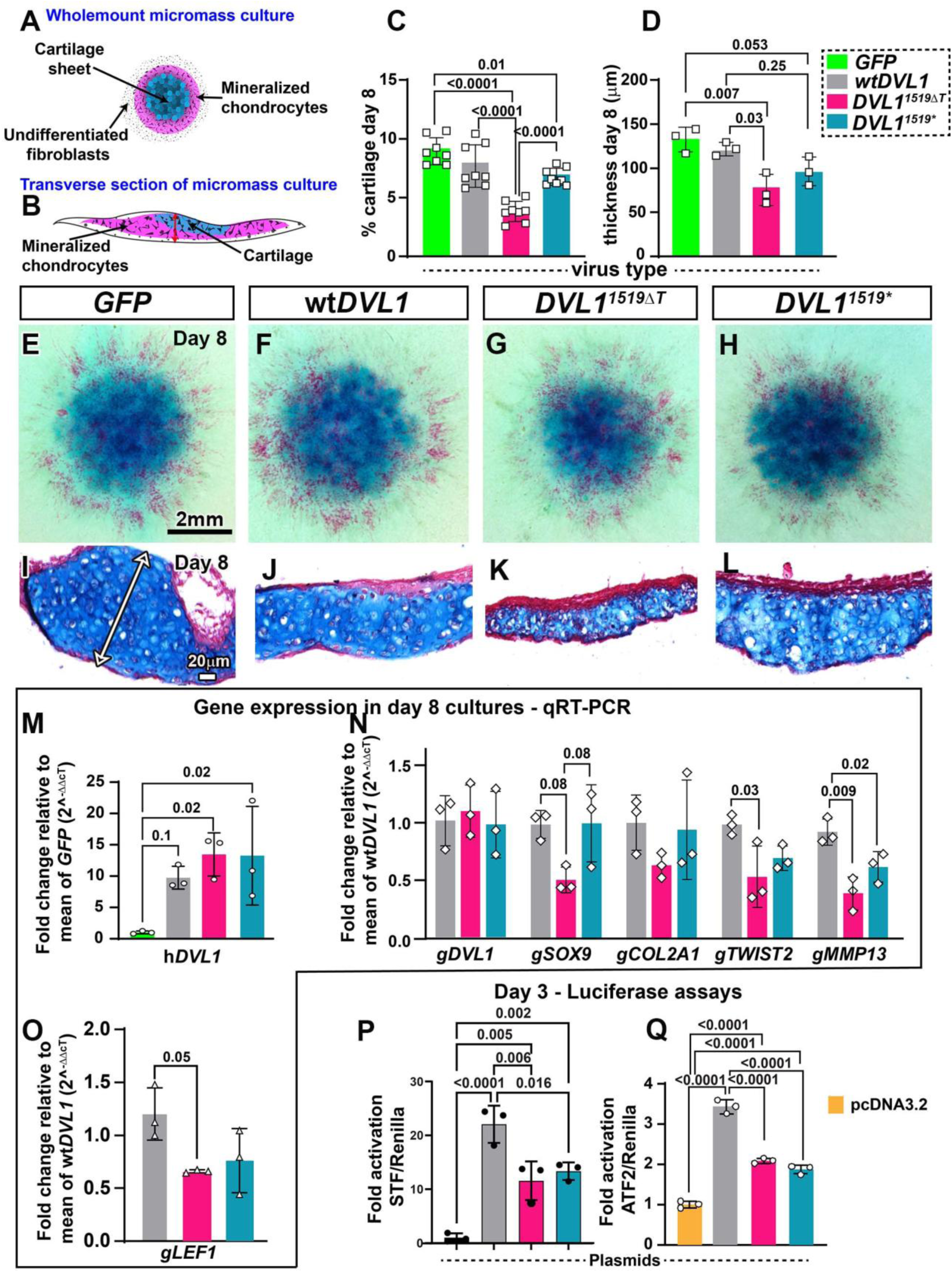
Effects of hDVL1 viruses on frontonasal mass micromass cultures: A) Mesenchymal cells harvested from the frontonasal mass of stage 24 (E4.5) embryos and were plated into high-density cultures. B) Other cultures were sectioned and used for microscopic analysis. C,E-H) There is a significant decrease in the proportion of cartilage from the DVL1^1519ΔT^ variant compared to the wtDVL1 infected cultures as measured in wholemount stained cultures. When the *DVL1^1519*^* truncation was compared to wtDVL1 there was no significant difference in the Alcian blue stained area. The *DVL1^1519*^* construct reduced cartilage compared to GFP controls. D,I-L) The cultures were significantly thinner in the presence of the *DVL1^1519ΔT^* variant compared to w*tDVL1* or the *GFP* controls. The *DVL1^1519^*\* construct slightly reduced the thickness of the cartilage compared to GFP controls. M) The viruses containing human DVL1 constructs were expressed at similar levels in primary mesenchyme as determined by qRT-PCR with human-specific DVL1 primers. Generally the human *DVL1* gene expression was elevated 10 to 15-fold by the viral transgenesis. N, O) The 1519ΔT virus significantly reduced expression of *TWIST2*, *MMP13* and *LEF1*. P,Q) Two reporters were used in micromass cultures from frontonasal mass cells, the SuperTOPFlash reporter for canonical WNT signaling and ATF2 for JNK-PCP, non-canonical WNT signaling. The wt*DVL1* plasmid significantly activated both reporters. In comparison both *1519ΔT* and *1519\** viruses did not activate the reporters as much as wt*DVL1*. They did retain more activity than the control, parent plasmid. Statistical analysis done with one-way ANOVA followed by Dunnett’s multiple comparison test (*M*) or Tukey’s post-hoc test (C,D,N,O,P,Q). Scale bar in E-H = 2 mm, I-L = 20µm.

Next, we quantified RNA levels of *SOX9* (early chondrocyte), *COL2A1* (cartilage matrix protein), *TWIST2* (cell differentiation), and *MMP13* (cleaves type II collagen) in day 8 micromass cultures (3 biological replicates with 12 cultures per replicate, Fig. 6N, Fig. S6). The levels of *SOX9* and *COL2A1* levels were significantly reduced in *DVL1^1519ΔT^* cultures compared to *GFP* cultures (Fig. S6) but the levels were comparable to wt*DVL1* and *DVL1^1519*^* cultures (Fig. 6N). The *DVL1^1519ΔT^* cultures also showed reduced levels of *TWIST2* RNA compared to those with *GFP* (Fig. S5) and wt*DVL1* cultures (Fig. 6N). These results are different than the immunostaining data for 96h specimens (Fig. 5I). It is possible that the appearance of increased TWIST protein reflects the location of both TWIST1 and TWIST2 since the immunogen shares 92% homology between the two proteins. We have reported that facial cultures exposed to WNT5A protein resorb the matrix due to induction of *MMP13* RNA (37). Interestingly, *MMP13* transcripts were significantly lower in *DVL1^1519ΔT^* and *DVL1^1519*^* cultures compared to wt*DVL1* cultures (Fig. 5N). Therefore, the presence of DVL1 variant is unlikely to contribute to cartilage matrix resorption. Indeed the level of Alcian blue staining is qualitatively similar in all cultures infected with the *DVL1^1519ΔT^* variant (Fig. 6G,K; S5C,G). This data on micromass cultures shows there is activity retained in the *DVL1^1519*^* gene in terms of inhibiting the continued secretion of cartilage matrix. Overall, decreased *SOX9*, *COL2A1*, and *TWIST2* in *DVL1^1519ΔT^* cultures suggests that the virus delays or inhibits matrix secretion thus reducing cartilage. DVL proteins play a vital role in the WNT pathway signaling (14). We observed lower levels of *LEF1* transcript (mediator of the WNT/βcatenin pathway) in *DVL1^1519ΔT^* cultures compared to wt*DVL1* (Fig. 6O). This suggests a signaling defect may also be present that needs to be tested further.

### *DVL1* C-terminus is crucial for both branches of WNT signaling pathway

We have previously shown that *DVL1^1519ΔT^* fails to activate the WNT/βcatenin pathway in HEK293T cells (7). To better preserve the signaling context we used frontonasal mass cultures to carry out canonical (42) and non-canonical luciferase assays (43). Similar to what we saw in HEK293T cells, *DVL1^1519ΔT^* weakly activated the SuperTOPFlash (STF) reporter and the ATF2 non-canonical WNT reporter (Fig. 6P,Q). Unexpectedly, *DVL1^1519*^* also exhibited weak activity (Fig. 6P, Q). Thus, our data suggest that in primary cells, the Robinow syndrome variant *DVL1^1519ΔT^* and the *DVL1^1519*^* truncation are both able to weakly activate the WNT canonical and JNK/PCP pathways. The presence of the full-length C-terminus in wtDVL1 is crucial to retaining full activity in both WNT signaling pathways.

### The abnormal DVL1 C-terminus increases JNK signalling in *Drosophila*

PCP defects in adult wing hairs can be caused both by loss or gain of Wnt/PCP pathway activity (44–46). Our previous work used two established reporters for to reveal the effect, the JNK targets *puckered-lacZ* (*puc-lacZ*) and Mmp1. Both reporters were used to show that DVL1 variants ectopically induced JNK signalling in the wing pouch of wing imaginal discs while *wtDVL1* did not show any induction (7). In this study, we did the same assays to see whether this induction is due to the lack of the original C-terminus or presence of the novel peptide. We used the transcriptional reporter *puc-lacZ* in larvae expressing the DVL1 proteins using the *apterous-Gal4* strain which leads to expression in the dorsal compartment of the wing disc (seen by Flag staining in Fig. 7B-D) at 29°C. Only the DVL1^1519ΔT^ expressing wing discs showed a robust induction of *puc-lacZ* signal (detected by anti-β-gal) in the dorsal compartment of the wing pouch (Fig. 7A-D’). We also used *dpp-Gal4* to express the DVL1 transgenes and quantified the expression of Mmp1. No induction of Mmp1 protein signal in controls, wt*DVL1* and *DVL1^1519*^* expressing wing imaginal discs were observed, while all the *DVL1^1519ΔT^* expressing discs showed significantly more Mmp1 puncta in the wing pouch within the *dpp-Gal4* expression domain (Fig. 7F-J).

**Figure 7.**
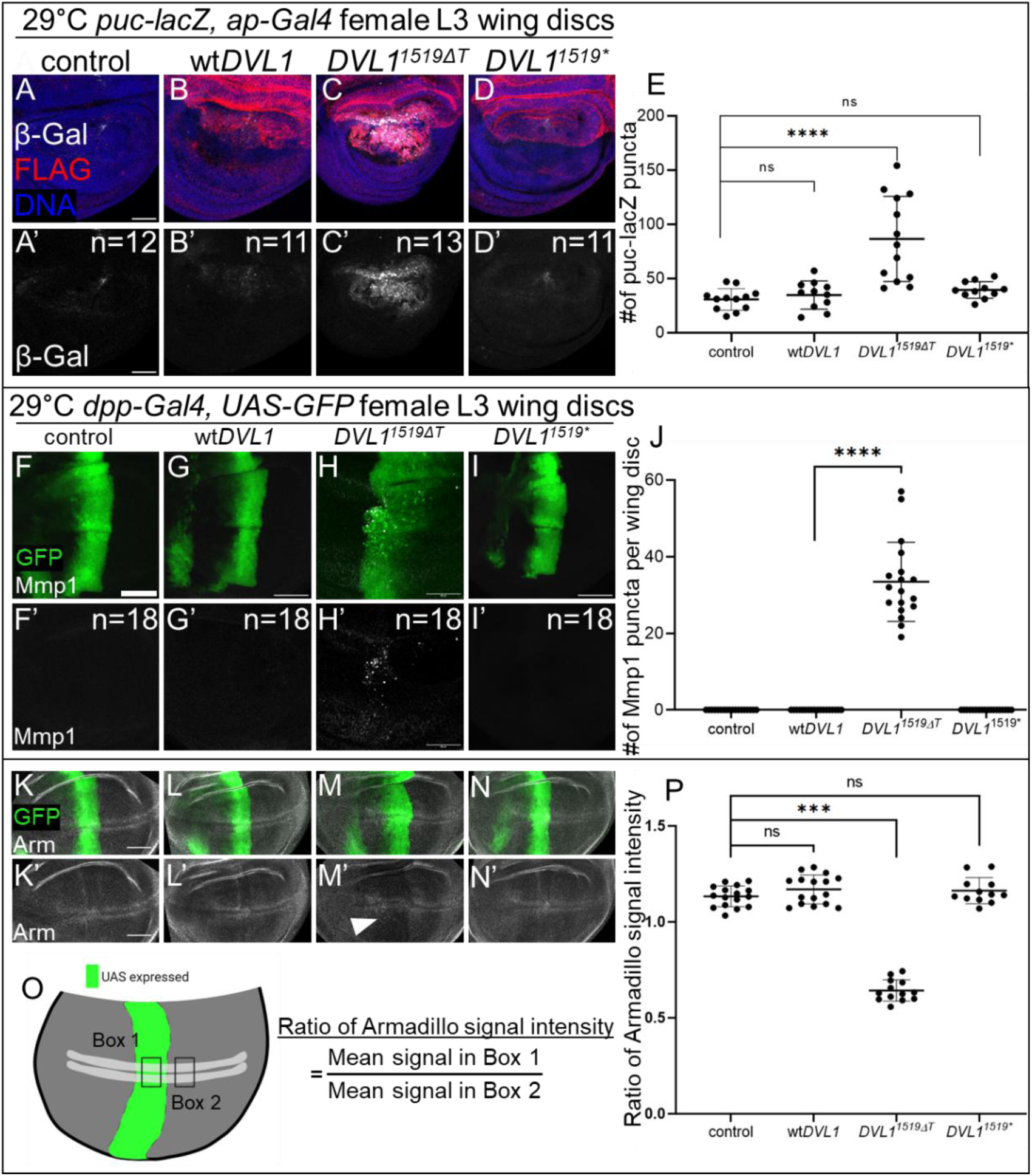
C-terminally truncated DVL1 proteins do not induce signaling defects in Wnt pathways in *Drosophila*. (**A-D**) Z-stack maximum projection of imaginal wing discs showing DNA (DAPI, blue), the expression of the genetically encoded Jnk reporter *puc-lacZ* (β-gal; white) and UAS transgene expression domain (red) staining in control (**A**) and *DVL1*-expressing female wing discs (**B-D**). (**A’-D’**) Z-stack maximum projection of single channel images showing β-gal (white) staining in control (**A’**) and *DVL1*-expressing female wing discs (**B’-D’**). n numbers in bottom panels depict the number of wing discs that displayed the phenotype shown in the representative image. Crosses were performed at 29°C and 11-14 female wing discs were imaged and quantified per genotype across n=3 independent experiments. Scale bar = 50 μm. (**E**) Plot of *puc-lacZ* puncta counted by finding signal intensity maxima in each wing disc imaged. Error bars show mean with SD. Statistics were performed with ANOVA. (**F-I**) Z-stack maximum projection of third instar wing imaginal discs showing Mmp1 (white) and *dpp>GFP* expression domain (green) in control (**F**) and *DVL1*-expressing female wing disc pouches (**H-I**). (**F’-I’**) Z-stack maximum projection of single channel images showing Mmp1 staining in control (**F’**) and *DVL1*-expressing female wing discs (**H’-I’**). Scale bar = 50 μm. (**J**) Plot showing the number of Mmp1 puncta counted per wing disc. Crosses were performed at 29°C and 18 female wing discs were analyzed per genotype. Statistics were performed with ANOVA. (**K-N**) Z-stack maximum projections of Arm protein (white) and dpp>GFP expression domain (green) staining in control (dpp>GFP,+) (**K**) and DVL1-expressing female wing discs (**L-N**) with the corresponding single channel Arm (white) staining shown in (**K’-N’**). Significant decrease in Arm levels relative to control is indicated with white arrow in M’. Scale bar = 50 μm. (**O**) Schematic of an imaginal wing disc pouch with dpp-Gal4 expression domain (green), position of most stabilized Arm protein (white). Boxes 1 and 2 show the regions where ratio of Arm signal intensity was quantified within and outside of the dpp expression domain. (**P**) The Arm signal intensity ratio quantified as described is plotted. Crosses were performed at 29°C and analyses show averages from 12-16 discs per genotype. Error bars show mean with SD. Statistics were performed with ANOVA.

### Lack of DVL1 C-terminus does not alter canonical Wnt/Wg signalling

In *Drosophila*, Armadillo (Arm) protein stability can be measured to investigate canonical Wnt signalling activity. Arm is the fruit fly ortholog of β-catenin, which is stabilized in the cytoplasm after Wnt signalling pathway activation. Arm expression can be seen as two stripes flanking the dorso-ventral boundary in the wing imaginal disc pouch. In our earlier work, it was shown that the expression of 3 Robinow syndrome *DVL1* variants caused a decrease in the Arm protein levels while the expression of *wtDVL1* did not have any such effects(7). We used *dpp-Gal4* to drive the expression of our transgenes at 29°C and measured the Arm signal intensity (Fig. 7O, P). We confirmed that only the *DVL1^1519ΔT^* variant altered the canonical/Wnt signalling while the C-terminal truncated protein *DVL1^1519*^*showed a similar phenotype to the control and *wtDVL1* (Fig. 7K-P). In the light of this finding, we concluded that the disruption in canonical Wnt signalling in *Drosophila* is not caused by the lack of the original C-terminus in Robinow syndrome-associated *DVL1* variant but the presence of the new peptide sequence.

### Variant *DVL1^1519^***^Δ^***^T^* protein accumulates within the nucleus whereas DVL1^1519*^ does not

The C-terminus of DVL is essential for protein-protein interactions (21, 47–50). In addition, the C-terminus has a nuclear localization sequence (NLS) and a nuclear export sequence (NES, M/LxxLxL) (Fig. 1A) that allows DVL shuttling and WNT signal transduction (22, 25, 51). The NES in *Drosophila* Dsh is located between amino acids 510-515 (22) but the exact location in human DVL1 is unknown. Based on the location of the frameshift in *DVL1^1519ΔT^* (p.TrpDVL1^507Glyfs*142^), we hypothesized that the *DVL1* variant lacks the NES, leading to abnormal subcellular localization. To eliminate effects of interactions with endogenous DVL1 or compensation from other family members, we overexpressed DVL1 plasmids in HEK293T DVL-triple knock out (TKO) cells, which have all three human *DVLs* knocked out (52). We determined that signal transduction in the canonical WNT pathway takes place when the wt*DVL1* is transfected into the TKO cells (Fig. S7A). However, neither the frameshifted variant nor the truncated version of DVL1 were able to activate the SuperTOPFlash reporter in this context unlike in the frontonasal mass cells (Fig. 6P). Due to the unavailability of DVL1 antibodies, we used Flag-tags to monitor DVL1 localization. The wt*DVL1* plasmid was distributed mainly in the cytoplasm (Fig. 8A-A’’; D,E). In contrast, the plasmid expressing Flag-tagged *DVL1^1519ΔT^* were primarily localized within the nucleus (Fig. 8B-B’’;D,E) while localization of plasmids encoding DVL1^1519*^ was similar to wt*DVL1* (Fig.8C-C’’,G,H,D,E). Both wt*DVL1* and *DVL1^1519*^*proteins showed characteristic cytoplasmic puncta which were almost completely absent in cells expressing *DVL1^1519ΔT^* (Fig.8A-C’’, arrowheads). We obtained similar nuclear distribution of the *DVL1^1519*^*variant protein in normal HEK293T cells (Fig. S7B). This suggests that novel C-terminus leads to mislocalization of protein coded by *DVL1^1519ΔT^* possibly due to interference of the novel C-terminus with the NES or changes in *DVL1* protein conformation.

**Figure 8.**
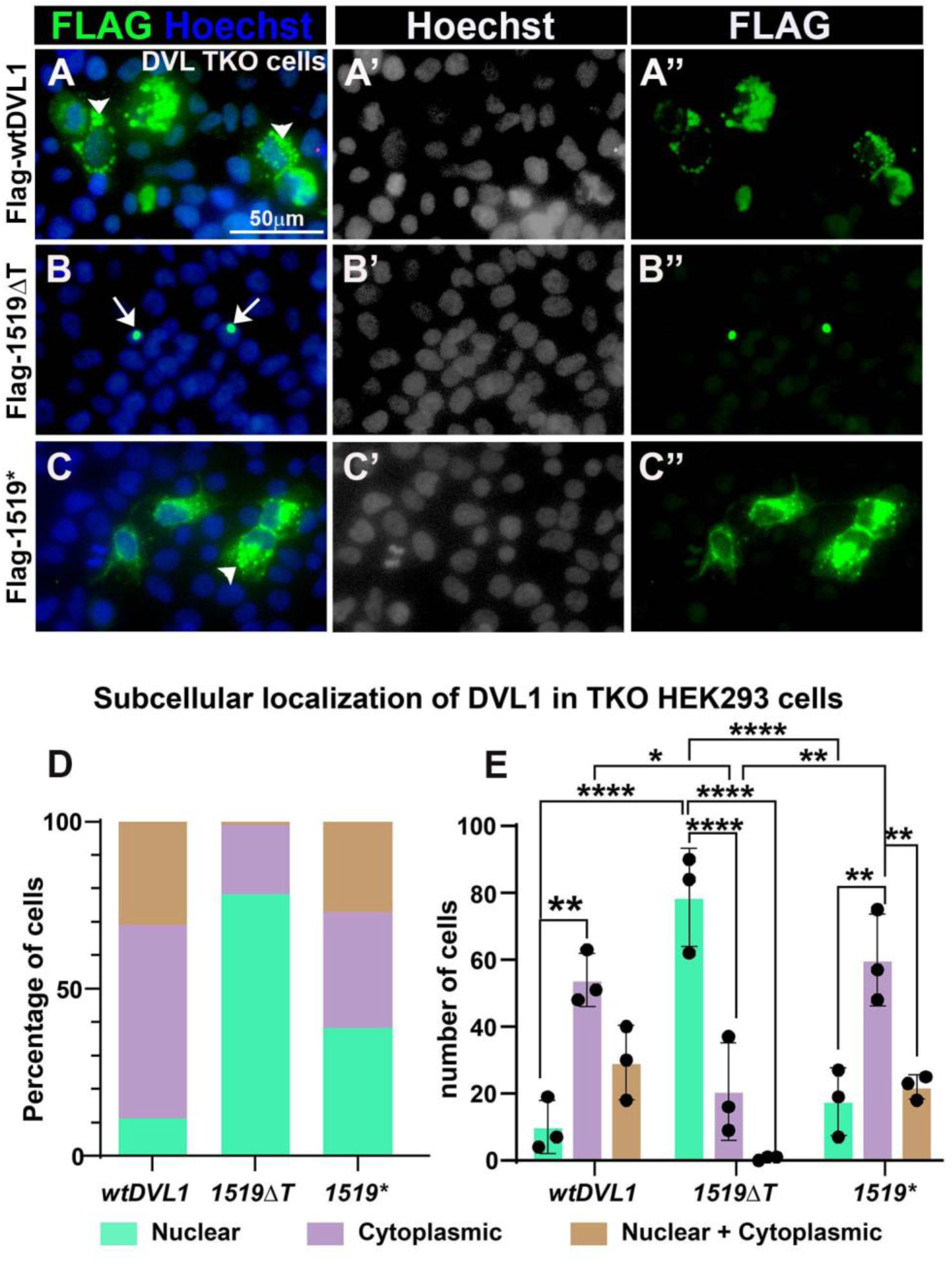
Plasmids expressing DVL1 variants expressed in triple knockout HEK293T cells lacking DVL1, 2 and 3. A-A’’) The expression of wtDVL1 as shown by the N-terminal Flag tag is mainly cytoplasmic. B-B’’) Small punctae of DVL^1519ΔT^ in the nuclei. C-C’’). In contrast the 1519* is expressed in the cytoplasm similar to wtDVL1. D,E) The proportion of cells expressing DVL^1519ΔT^ in the nuclei is significantly higher than for wtDVL1 and 1519*. Statistical analysis done with 2-way ANOVA followed by Tukey’s post-hoc test (E). Scale bar = 50µm for all panels. P values *= p<0.05, **= p< 0.01, *** =P<0.001, ****=P<0.0001.

## Discussion

The discoveries made in this work include the first insights into the mechanism for wide set eyes and broad nasal bridge seen in patients with all forms of Robinow syndrome. We also tested the 1519* truncated version of the protein to separate the requirements for the wildtype C terminus versus neomorphic effects of the abnormal C terminus. The mystery surrounding the abnormal C terminus that is present in every *DVL1, DVL2* and *DVL3* mutation associated with autosomal dominant Robinow syndrome led some researchers to speculate that this causes a dominant interference with the normal protein (12, 13). The novel C-terminal peptide has no known homology and is conserved among individuals with the *DVL1* form of Robinow syndrome (1, 13). In individuals with *DVL3* frameshift mutations, a different extended C-terminal peptide is generated (7) so the specific sequence is not contributing to the phenotype. Our study uses a truncated version of *DVL1* that has shed light on the requirements for the normal and abnormal *DVL1* C-terminus in development, skeletal differentiation and cell signaling.

### Connections between the *DVL1* frameshifts, chicken facial morphogenesis and the facial phenotypes in Robinow syndrome

All Robinow syndrome individuals have a wider nasal bridge, hypertelorism, frontal bossing and maxillary hypoplasia (9, 11, 53). Some of the phenotypes produced in chicken embryos can explain the origins of these specific Robinow syndrome dysmorphologies. We found that the frontonasal mass width was increased by the h*DVL1* variant. These effects are unrelated to neural crest cell migration since viruses are injected after neural crest cells have settled into the face. We also ruled out increased proliferation as the mechanism contributing to facial widening since proliferation is unchanged at 48h and then decreases by 96h. We had also shown that over expression of two human *FZD2* variants caused a relative increase in the width of the frontonasal mass by 72h without changing proliferation (8). We also showed that overexpression of human *DVL1* variants had no effect on endogenous genes compared to wt*DVL1*. We suggest that the variants in DVL1 and FZD2 prevent the normal process of narrowing of the frontonasal mass (41). The mechanism of narrowing is related to cytoskeletal arrangement regulated by the RhoA-GTPase pathway. In a previous study from our lab where ROCK (the direct target of RhoA) was inhibited, the frontonasal mass failed to narrow (41). Since h*DVL1* signals via DAAM to activate the RhoA and Rock pathway (54), it is possible that this branch of WNT signaling is less active in the presence of the variant form of h*DVL1*. Indeed, ATF2 luciferase was less active in the presence of *DVL1^1519ΔT^* compared to wt*DVL1*. We also noted that the midfacial cartilage is not patterned correctly and therefore the narrow nasal prenasal cartilage (equivalent of the nasal septum) fails to form. This disruption would be sufficient to cause the broad nasal bridge and shorter midface. We also noted a decrease in chondrogenesis in micromass culture so a failure of cartilage differentiation in the human midline could contribute to less midfacial outgrowth. The nasal septum is a major growth site for the midface (55).

Other facial phenotypes produced by the h*DVL1* overexpression in chicken embryos such as the shorter beak and curvature towards the injected side recapitulates midface hypoplasia observed in patients with autosomal dominant Robinow syndrome (9, 12). Indeed, the h*DVL1^1519ΔT^* variant caused significant shortening and deviation of the upper beak with a 92% penetrance. It is possible that the decreased proliferation we identified in stage 29 embryos could explain the decreased length of the upper beak. Other mechanisms underlying jaw deficiencies seen in Robinow syndrome could be a failure of membranous bone differentiation as we reported for the FZD2 variants (8). The facial bones such as premaxilla, maxilla and mandible are formed by direct ossification of the neural crest-derived mesenchyme (56). There is no cartilage precursor for intramembranous bone, therefore the jaw hypoplasia in Robinow syndrome patients is due either to smaller condensations in the early stages or decreased apposition of bone in later stages of bone remodeling. In our study the premaxilla failed to form in 12/13 embryos injected with *DVL1^1519ΔT^* virus. This suggests that the *DVL1^1519ΔT^* variant interferes with the early mesenchymal condensation stages whereas the FZD2 variants interfere with later stages of mineralization since all bones are present in these animals (8).

### The effects of *DVL1* on canonical WNT signaling vary according to context

We showed that the *DVL1^1519ΔT^* variant was significantly less able to activate the canonical luciferase reporter (STF) in primary facial mesenchyme compared to wt*DVL1*. The h*DVL1^1519ΔT^* variant was transfected into HEK293 cells by our group in a previous study and a similar decrease in STF signaling was observed (7). In that study we also showed that the equimolar combination of h*DVL1^1519ΔT^* and wt h*DVL1* led to a dominant-negative inhibition of the wt protein in luciferase assays. The substitution of the abnormal C-terminal peptide removes an important binding site for a protein that is needed for canonical WNT signal transduction. For example, IQGAP1 (Ras GTPase-activating-like protein) associates with the C-terminus of DVL1 and has been shown be essential to activate β-catenin mediated canonical WNT activity (49). We hypothesize that the loss of activity in the canonical pathway may be attributed loss of interaction of variant h*DVL1* with a protein such as IQGAP1.

Nuclear localization of β-catenin and displacement of the Groucho co-repressor is required for increased transcriptional activity in the canonical WNT pathway (4, 57). We expected to see reduced nuclear β-catenin in vivo in the frontonasal mass presence of variant *DVL1* but there were no changes in the proportion of positive cells. The levels of total g*CTNNB1* were measured 48h after virus infection in vivo. The RNA levels of g*CTNNB1* were also not changed. The qRT-PCR experiments of micromass culture collected at day 8 also showed no significant changes in g*CTNNB1* with the autosomal dominant Robinow syndrome variant. Taken together our data suggests the h*DVL1^1519ΔT^* transiently inhibits canonical WNT signaling during critical stages when intramembranous bone condensations are developing.

### What did we learn about normal C-terminus function and changes that occur due to the Robinow syndrome frame shifts?

One of the most unexpected findings was that expression of a human DVL1 construct lacking almost all of the C-terminus (*DVL1^1519*^*codes for DVL1^507*^) permits normal development in chickens and flies. Several key functions are lost in this shorter form on DVL1. The looping back of the C-terminus to the PDZ domain does not occur, therefore there will be lower levels of JNK signaling (20, 22, 58). We did measure less activity in the ATF2 reporter when frontonasal mass cells were transfected with *DVL1^1519*^*. However, the main regulation of JNK signaling depends on the binding of DVL1 via the DEP domain to FZD. These levels of activity must be sufficient to support development (58).

Protein degradation would be reduced due to loss of E3 ligase binding sites in the shorted form of DVL1. Indeed, in western blots of human gene constructs expressed in Drosophila there is significantly more protein derived from *DVL1^1519*^* compared to the *DVL1^1519ΔT^* construct. Therefore, either the higher levels compensate for the loss of the C-terminus conformation change or that variation in protein levels are not critical for developmental functions. Even though there is less protein degradation, the nuclear and cytoplasmic distribution of DVL1^507*^ in HEK293 TKO cells is indistinguishable from wtDVL1. This data suggests that nuclear export is occurring normally and that the NES of DVL1 is localized to the first 63 amino acids of the C terminus.

In contrast, the *DVL1^1519ΔT^* variant had major effects on development and signaling in all assays. In the PCP signaling JNK pathway in *Drosophila* only the 1519ΔT variant can stimulate the readouts suggesting there is a dominant impact. Similarly, the 1519ΔT variant causes a repression of canonical WNT signaling in *Drosophila.* These data are different to luciferase assays carried out in frontonasal mass cells where weak activation of the ATF2 reporter was seen. Thus, the vertebrate may respond to the *DVL1^1519ΔT^* variant differently than in the fly. The morphogenesis of the fly and chicken were both severely disrupted by *DVL1^1519ΔT^*, again showing that the abnormal C terminus is interfering with normal DVL1 or Dsh function. In addition to signaling defects, other molecular mechanism could involve pleiotropic effects on many other signaling pathways since there is nuclear translocation of the variant protein. Finally, the conformational changes of the abnormal DVL1 protein have not been determined and the frame-shifted C terminus likely has low predictive value in terms of tertiary structure. Indeed the full predicted structure of DVL protein is still not known due to the 3 intrinsically disordered domains and various post-translational modifications (58). These authors predicted the DVL3 3D conformation using AlphaFold3 and focused on the DIX, PDZ and DEP domain rather than the C terminus (58).

Of the three forms of DVL, DVL2 makes up 80-95% of the protein in cells (59). Therefore, it is interesting that variants of such a low abundance protein as DVL1 can impact development. The presence of DVL2 and DVL3 cannot compensate for the abnormal C terminus of DVL1. A similar proposed dominant mechanism likely applies to DVL2 (1) and DVL3 (1, 18) forms of Robinow syndrome. Our findings support these original hypotheses put forward by those that first identified the *DVL1* variants that cause Robinow syndrome.

## Materials and methods

### Chicken embryo model

White leghorn eggs (*Gallus Gallus*) obtained from the University of Alberta were incubated to the appropriate embryonic stages, based on the Hamilton Hamburger staging guide (60). All experiments were performed on prehatching chicken embryos which are exempted from ethical approval by the University of British Columbia Animal Care Committee and the Canadian Council on Animal Care.

### Cloning of human wt and variant human *DVL1* constructs

The open reading frame encoding human DVL1 was purchased from Origene (#RC217691). The insert was initially in a pCMV6 vector containing a full coding sequence with two C-terminal tags: myc and DDK (Flag similar). To change pCMV6 vector to an entry clone (pDONR221), gateway cloning with BP clonase II (ThermoFisher #11789020) was performed. Then restriction free cloning was used to insert kozak- N-terminal flag (DYKDDDDK) sequence and 3’ C-terminal STOP (added to eliminate commercial tags from company) to this pDONR221 vector. Site-directed mutagenesis was performed in the pDONR221 (Gateway compatible) vector and then recombined into destination vectors (pcDNA3.2 expression vector and RCAS retroviral vector) using LR Clonase II enzyme (ThermoFisher # 11791019). Gateway cloning (ThermoFisher) was used to move human *DVL1* into the pDONR221. Site-directed mutagenesis was used to knock-in the mutations. Autosomal dominant Robinow syndrome *DVL1* frameshift mutation (OMIM: 616331) was knocked in (*1519ΔT*) (White et al., 2015). One additional h*DVL1* construct was made for this study - A STOP codon was placed immediately after nucleotide 1519 (*DVL1^1519*^* coding for DVL1^507*^) as published by others (12). To create the RCAS viruses, a Gateway compatible RCASBPY destination vector was used for recombination with the pDONR221 vector using LR Clonase II (61). The location of *DVL1* containing viruses was determined retrospectively on histological sections stained with a Group-associated antigens (GAG), an antibody that recognizes specific proteins in the RCAS virus. To generate the plasmids for luciferase assays and immunocytochemistry, the inserts were cloned into the Gateway compatible pcDNA3.1 vector.

### Growth of RCAS viral particles and viral titre

Replication-Competent ASLV long terminal repeat (LTR) with a Splice acceptor (RCAS) (61, 62) plasmid DNAs (2.5µg) encoding *GFP*, human *DVL1*, or three *DVL1* variants were transfected into the DF-1 immortalized chicken fibroblast cell line (American Type Culture Collection, CRL-12203) using Lipofectamine 3000 (Thermo Fisher #L3000-008) following the manufacturer’s guidelines. RCAS virus containing *GFP* insert (kindly provided by A. Gaunt) served as a control in virus overexpression studies, as published(7, 34, 35, 37, 63). DF1 cells were cultured at 37°C and 5% CO_2_ in DMEM (ThermoFisher#1967497) medium supplemented with 10% fetal bovine serum (FBS; Sigma #F1051) and 1% penicillin/streptomycin (ThermoFisher #15070-063). The cells were maintained in 100 mm culture dishes with media changes every other day and passaged 1:2 two to three times per week using trypsin-EDTA (0.25%, ThermoFisher #25200-072). After six weeks of culturing, the viral particles were collected and centrifuged in a swing bucket SW28 rotor (Beckman #97U 9661 ultracentrifuge) for 2.5 hours (no brake) at 25,000 rpm at 4°C. The supernatant was carefully removed, and the resulting pellet was resuspended in 50-100 µl Opti-MEM (ThermoFisher#319850962). This suspension was then incubated overnight at 4°C. The concentrated viral particles obtained were aliquoted (5µL aliquots), rapidly frozen in methanol + dry ice, and stored at −80°C for future use (64).

To determine viral titer, 50-60% confluent DF1 fibroblasts were infected with serial dilutions of 2μl of concentrated viral stock. After 36 hours of virus incubation, cells were fixed in 4% paraformaldehyde for 30 mins. Immunocytochemistry with Group Associated Antigens (GAG) antibody (Developmental Studies Hybridoma bank, AMV-3C2) was performed on virus-treated cells (Table S7). The cells were permeabilized for 30 mins with 0.1% Triton X-100, followed by blocking in 10% goat serum and 0.1% Triton X-100 and overnight incubation with the primary antibody (Table S7). Fluorescence images were captured using a Leica inverted microscope at 10x with a DFC7000 camera. The analysis of viral titer was done with ImageJ’s cell counter tool by determining the proportion of GAG-positive cells per mL of virus plated in a 35mm culture plate. Virus titer = # of GAG positive cells * (Area counted * total cells expressing GAG)/2*1000 (Fig. S7).

### Chicken embryo injections

Fertilized eggs obtained from the University of Alberta, Edmonton, were incubated in a humified incubator at 38°C until Hamilton Hamburger (60, 65) stage 15 (E2.5). Concentrated RCAS (titer = >2 x 10^8^ IU/mL) retrovirus viral particles (∼5μl) combined with Fast Green FCF stain (0.42%, Sigma #F7252) (1μl) were injected into the frontonasal mass (anatomic region bounded by the nasal slits) of stage 14-15 chicken embryos (25-28 somites) using glass filament needles (thin-wall borosilicate capillary glass with microfilament, A-M systems #615000) and a Picospritzer microinjector (General valve corp. #42311). The infection of embryos with RCAS at stage 15 (E2.5) was performed to ensure maximum infection of facial prominences(34, 63). Due to accessibility, all injections were made into the right frontonasal mass as the chick embryos turn on their left side during development. The facial prominences form around stage 20, and the complex and temporally regulated patterning occurs between stages 20-29. The skeletal derivatives of the frontonasal mass are fully patterned and ossified between stages 34-40. The investigation encompassed multiple embryonic stages to comprehensively analyze these developmental processes. After conducting the overexpression of high-titer *DVL1* viruses at stage 15, the retrospective determination of virus location was carried out on histological sections. These sections were stained with Group-associated antigens (GAG), an antibody that identifies specific proteins in the RCAS virus (7) (Table S7).

### Wholemount staining of skulls

To study skeletal elements, embryos were grown until stage 38 (10 days post-injection, Table S3). The embryos were washed in 1x phosphate-buffered saline (PBS; 137 mM NaCl, 8.1 mM Na_2_HPO_4_, 2.7 mM KCl, 1.5 mM KH_2_PO_4_; pH 7.3) and fixed in 100% ethanol for 4 days. After removal of eyes and skin, the embryos were transferred to 100% acetone for another 4 days. Subsequently, the heads were stained with a freshly prepared bone and cartilage stain (0.3% Alcian blue 8GX (Sigma #A5268) in 70% ethanol, and 0.1% alizarin red S (Sigma # A5533) in 95% ethanol, with 1 volume of 100% acetic acid and 17 volumes of 70% ethanol) for two weeks on shaker at room temperature. Following staining, the skulls were washed in water and cleared in a 2% KOH/20% glycerol solution on shaker for 4 days, followed by immersion in 50% glycerol for imaging. The heads were stored in 100% glycerol post-imaging. Phenotyping was conducted by photographing the right lateral, superior and palatal views of cleared heads using a Leica DFC7000T microscope camera. Skeletal preparations from each virus type were analyzed for changes in the size or shape of bones derived from the frontonasal mass, missing bones, or qualitative reduction in ossification observed as reduced alizarin red stain. Statistical analysis of the frequency of observations was performed using contingency analysis with Fisher’s exact test in GraphPad Prism 10.1.0.

### Primary cultures of frontonasal mass mesenchyme

Stage 24 chicken embryos were extracted from the eggs and their extra-embryonic tissues were removed in cold phosphate-buffered saline (PBS). The frontonasal mass was dissected in cold Hank’s balanced saline solution (HBSS) (without calcium and magnesium) (ThermoFisher #14185052) with 10% FBS and 1% Antibiotic-Antimycotic (Life Technologies #15240-062). Dissected frontonasal pieces were incubated in 2% trypsin (Gibco) at 4°C for 1 hour. Hank’s solution was added to inhibit the enzymatic activity of trypsin. Ectoderm was manually peeled off from the frontonasal mass pieces. The cell solution was then centrifuged at 1000 g, 4°C for 5 minutes. The supernatant was removed and the frontonasal mass pieces were resuspended in Hank’s solution. The mesenchymal cells were counted using a hemocytometer and 2x107 cells/ml were resuspended in chondrogenic media (micromass media) containing DMEM/F12 medium (Corning #10-092-CV) supplemented with 10% FBS, 1% L-Glutamine (ThermoFisher #25030), Ascorbic acid (50 mg/ml) (ThermoFisher #850-3080IM), 10 mM β-glycerol phosphate (Sigma Aldrich #G9422) and 1%, Antibiotic-antimycotic (ThermoFisher #15240-062). The cells in suspension were subsequently infected with 3µL *GFP* (control), wt h*DVL1*, or h*DVL1* variants containing viruses. The 10µl of cells suspension infected with virus was plate micromass cultures (3-4 spots per 35mm culture dish, NUNC #150318) at a density of 2 x 10^7^ cells/ml, (37, 66). The culture plates were incubated at 37 °C and 5% CO_2_ for 90 minutes to allow cells to attach and then flooded with 2 ml of micromass media. Thereafter, micromass culture media was changed every other day for experimental time points of day 4, 6, and 8.

### Wholemount staining of micromass cultures

On day 4, 6, and 8, cultures were fixed in 4% paraformaldehyde for 30 minutes at room temperature and subjected to wholemount staining. To detect the mineralization of cartilage using alkaline phosphatase stain (Table S8), fixed cultures were incubated at room temperature in 100mM Tris for 30 minutes (pH 8.3). Following this, the cultures were stained with 0.5% Alcian Blue in 95% EtOH: 0.1 M HCl (1:4) to detect the area occupied by cartilage, as previously described (Hosseini-Farahabadi, 2013; Underhill, 2014). All cultures were counter stained with 50% Shandon’s Instant Hematoxylin (ThermoFisher #6765015). The stained cultures were photographed under standard illumination using a stereomicroscope (Leica #M125). Wholemount staining was conducted on three biological and three technical replicates, and the experiment was repeated five times.

### Histology and Immunofluorescence

Embryos collected at stage 24 or 29 (Table S3) or micromass cultures (day 4, 6, 8, Table S5) were fixed in 4% PFA. The embryo samples were immersed in the fixative for 2-3 days at 4°C. The RCAS-infected cultures fixed in 4% paraformaldehyde for 30 mins. The cultures were then removed from the plate using a cell scraper (ThermoFisher #08-100-241), embedded in 2% agarose (Sigma #A9539) on a cold ice slab, and subsequently wax-embedded. The embryos (positioned frontally) and cultures (positioned transversely) were embedded in paraffin wax and sliced into 7µm sections. The sections were then utilized for histological and immunostaining analysis.

Selected frontal (embryos) and transverse (micromass cultures) sections were stained to visualize the differentiated cartilage and bone. Sections were dewaxed in xylene, rehydrated from 100% ethanol to water, and stained with 1% Alcian blue 8GX (in 1% acetic acid) for 30 minutes. After staining, sections were rinsed in 1% acetic acid and water. Subsequently, sections were stained in Picrosirius Red (0.1% Sirius Red F3B in saturated picric acid) for 1 hour in dark, followed by rinsing in 1% acetic acid, dehydration through ethanol, back to xylene, and mounted with Shandon Consul-mount (Thermo Scientific #9990441).

Immunofluorescence analysis was conducted on in vivo and day 6 and 8 cultures. Specific antibodies and treatments performed for each assay are outlined in Table S7. Primary antibodies were allowed to incubate overnight at 4°C, while secondary antibodies were incubated at room temperature for 1.5 hours unless otherwise specified. Sections were counter stained with Hoechst (10μg/ml #33568, Sigma) and incubated for 30 minutes at room temperature, then mounted with Prolong Gold antifade (ThermoFisher #P36930). Fluorescence images were captured using 20X objective on a slide scanner (3DHISTECH Ltd., Budapest, Hungary).

### Apoptosis and Cell proliferation

Apoptosis was analyzed using TUNEL (Terminal deoxynucleotidyl transferase dUTP nick end labeling) assay on sections obtained from virus infected frontonasal mass stage28 and micromass cultures sections day 6 and day 8. The TUNEL assay was performed using ApopTag Plus *in Situ* Apoptosis Fluorescein Detection Kit (Millipore Sigma # S7111).

For cell proliferation studies, embryos at stage 28, stage 29 or stage 30 were labeled with 50µl of 10 mM BrdU (Bromodeoxyuridine; Sigma #B5002) and incubated at 38°C for 1 hour prior to euthanizing. For labeling micromass cultures, 50µl of 10mM BrdU was added to the culture media (37°C 5% CO_2_) 1 hour before fixing day 6 and day 8 cultures in 4% PFA. Immunostaining was performed on the sections with anti-BrdU (Developmental Studies Hybridoma bank, 1:20, #G3G4) as described in Table S7. Fluorescence images were collected with a 20X objective on a slide scanner (3DHISTECH Ltd., Budapest, Hungary).

### Immunocytochemistry with transfected human *DVL1* variants

HEK293T DVL Triple Knockout (TKO) cells were received as a kind gift from Dr. Stephane Angers (University of Toronto) (52). Cells were grown to 60% confluency and transfected using Lipofectamine3000 (ThermoFisher #L3000-008) following manufacturer’s instructions. Transfected plasmids include (2.5 μg DNA): parent plasmid pcDNA3.2, wt*DVL1, DVL1^1519ΔT^, DVL1^1519*^*. Post transfection, the cells were grown for 48h and then fixed in 4% PFA for 20 min and then stored in 1X PBS at 4°C overnight. Cultures were blocked in 10% normal goat serum with 0.2% triton-X in 1 X PBS for 1h then incubated overnight at 4°C with primary anti-Flag antibody. Secondary antibody was applied for 1h at room temperature and counter stained with 10 µg/ml Hoechst (Table S8). Cells were imaged using 20x dry objective Leica THUNDER 3D cell imager widefield microscope with a Leica K8 camera.

### qRT-PCR on frontonasal mass in vivo and in vitro

Viral spread in the frontonasal mass was quantified using primers specific to human *DVL1* (primer set, Table S6). Three biological replicates containing five to six pieces of the right half of the frontonasal mass pooled in each sample were harvested for each virus at 48 h post injection. Similarly, three biological replicates containing pools of 12 micromass cultures per replicate were collected on day 6 and day 8. Total RNA was isolated from frontonasal masses using Qiagen RNAeasy kit (#75144, Toronto, Canada). Sybr green-based quantitive reverse transcriptase polymerase change reaction (Advanced universal SYBR® Green supermix; Bio-Rad #1725271) (qRT-PCR) was carried out using an Applied Biosystems StepOnePlus instrument. qRT-PCR cycling conditions were - 95°C for 10min, 40X (95°C for 5s, 60°C for 20 seconds). Analysis used human-specific primes for *DVL1* and avian primers were used (Table S6). The expression of each biological replicate was normalized to 18s RNA (Applied Biosystems, 4328839) and then these ΔCt values were used to calculate the ΔΔCt relative to the average levels of expression of the gene in *GFP*-infected cultures. The ΔΔCt method was used to calculate relative fold-change expression between Robinow syndrome-*DVL1* infected frontonasal mass and *GFP* as described (67). Statistical analysis was done with one-way ANOVA Tukey’s post hoc test in GraphPad Prism 10.0.2. A sample size calculator was used to determine how many samples would need to be included in order to detect a P value of 0.05 80% of the time and it was necessary to collect 13 biological replicates. This number of biological replicates was not feasible for these studies.

### Luciferase reporter assays

Transient transfections for luciferase assays were performed in untreated stage 24 frontonasal mass mesenchymal micromass cultures (37, 63). Cells were transfected with Lipofectamine 3000 (Invitrogen, L3000-008, Nunc 24-well plates #142475) transfection reagent. Micromass cultures were allowed to attach for 45 mins after plating and transfection reagents were added to the culture spot 30 mins prior to flooding the culture plate with micromass media. The following plasmids were used: control/empty (pcDNA3.2/V5-DEST), h*DVL1*, *DVL1^1519ΔT^*, *DVL1^1519*^*, *DVL1^1431*^*. Firefly reporter plasmids: SuperTOPFlash (STF; 0.2ug, Addgene plasmid #12456) and Activating Transcription Factor 2 (0.4ug; ATF2) (43) along with Renilla luciferase was transfected for normalization (0.01μg). Renilla luciferase was used as normalization control. Assay reading was done 48h after transfection representing day 3 of culture. The dual-luciferase reporter assay system (Promega #E1910) was used for all luciferase assays as described (63). Luminescence activity was detected with a PerkinElmer Victor X2 Multilabel Microplate Reader at 1 s reading with OD1 filter. All data shown represents two to three independent experiments with three technical and three biological replicates carried out for each transfection mixture. Statistical analysis done using one-way ANOVA, Tukey’s post hoc test in GraphPad Prism 10.0.2. The number of biological replicates was determined by our previous studies using luciferase assays (7, 35).

### Image analysis and statistics

For measurement of the width of the frontonasal mass at stage 29, distance between the nasal slits (illustrated in Fig. 5R) was measured manually with linear measurement annotation tool in CaseViewer (version 2.4). The data was analyzed using one-way ANOVA, Tukey’s test. The thickness of day 8 micromass cultures was measured using histological sections stained with Alcian blue and picrosirius red. The linear measurement annotation tool in CaseViewer (v2.4) was utilized. For each genotype, measurements were performed on three technical replicates from adjacent sections that were averaged to give one biological replicate. There were 3 biological replicates per genotype. The data was analyzed using one-way ANOVA, Tukey’s test in GraphPad Prism 10.1.0.

For analysis of immunofluorescence staining performed on stage 24 and 29 samples there were between 3-6 specimens analyzed (Table S3). For BrdU quantification, the right frontonasal mass was divided into four regions (100 x 250um^2^) (Fig. 4M, 5P) to count the proportion of cells expressing BrdU (N=4 for stage 24 and N = 3 for stage 29) and β-catenin (n=3 for stage 29). All cell counts were performed twice with the counter plugin in ImageJ by a blind observer. The data was analyzed using one-way ANOVA, Tukey’s test in GraphPad Prism 10.1.0. Similar samples sizes were used in other studies on BrdU labelling (7, 35).

For analysis of subcellular location of Flag-DVL1 protein in HEK293T DVL TKO cells, 150 transfected cells in random fields of view were counted in three biological replicates (three culture plates) (450 cells per virus type). The expression of FLAG was classified as cytoplasmic, nuclear, cytoplasmic + nuclear, and DVL1 forming puncta. Statistical analysis was conducted using two-way ANOVA, Tukey’s post hoc test in GraphPad Prism 10.1.0.

### Drosophila Husbandry

Flies were raised on standard media. Stocks were kept at room temperature (∼22°C) and crosses were performed at 25°C or 29°C as indicated. Three Gal4 driver fly lines were used to induce transgene expression: *dpp-Gal4, UAS-GFP*/TM6B (Swarup, Pradhan-Sundd and Verheyen, 2015), *Dll-lacZ/Cyo; Hh-Gal4/TM6B* (Hall *et al.*, 2017)*, ap-Gal4, UAS-GFP;+/SM6a∼TM6B*. Additional stocks used were: *puc-LacZ* (Martin-Blanco *et al.*, 1998)*, w^1118^* (RRID:BDSC_5905), *UAS-GFP*, *UAS-myr-RFP* (RRID:BDSC_7118).

### Dissections

#### Wing imaginal discs

Third instar larval wing imaginal discs were dissected out of larvae in cold PBS. The top half of the larvae were separated and flipped inside out to expose the imaginal discs. Fat tissue and the guts were removed while leaving the imaginal tissue attached to the cuticle.

#### Pharate Pupae

Pupae were gently pulled off the sides if vials and placed on double sided tape on a microscope slides. Forceps were used to pry open the operculum and tear away the case. Pharates were then photographed using a Leica dissection microscope and an iPhone.

#### Adult wings

Adult flies of the target genotype were collected in small glass vials filled with 70% ethanol. The wings were dissected using forceps in 95% ethanol in dissection dishes and mounted immediately. Aquatex (EMD Chemicals) was used as mounting media. The slides were baked overnight at 65°C incubator with small weights on them.

### Immunofluorescence staining, microscopy, and image processing

Tissue was dissected in phosphate-buffered saline (PBS) and fixed in 4% PFA at room temperature for 15 minutes. Samples were washed twice for 10 minutes with PBS with 0.1% Triton X-100 (PBS-T). Following a 1-hour block with 5% bovine serum albumin diluted in PBST at room temperature, samples were incubated overnight with primary antibodies at 4°C. Antibodies are listed in Table S8.

Samples were washed twice for 10 minutes with PBS-T (0.5% Triton-X100 in PBS) and incubated with Cy3 and/or Alexa Fluor 647-conjugated secondary antibody (1:500, Jackson ImmunoResearch Laboratories), DAPI (4’,6-Diamidino-2-Phenylindole) and/or Rhodamine Phalloidin (Invitrogen™ R415). for 2 hr at room temperature. After two 10- minute washes, samples were mounted in 70% Glycerol in PBS or VECTASHIELD® Antifade Mounting Medium (Vector Labs) and imaged using a Nikon Air laser-scanning confocal microscope or a Zeiss LSM880 with Airyscan confocal microscope. Images were processed with ImageJ, FIJI software (Schindelin *et al.*, 2012; Schneider, Rasband and Eliceiri, 2012) and are presented as Z-stack maximum intensity projections unless otherwise stated.

### Quantification of *puc-lacZ* signal intensity in wing discs

After completing the staining and mounting steps, wing imaginal discs were imaged as described previously. The signal intensity was measured by using the measurement tools in ImageJ software (Schindelin *et al.*, 2012): Using a rectangular box of identical dimensions, mean β-Galactosidase signal intensity was quantified inside and outside (dorsal and ventral compartments) of the transgene expression domain. The ratio of signal inside/outside β-Galactosidase signal was used to determine *puc-lacZ* levels within the transgene expression domain.

### Quantification of Armadillo protein levels

Imaginal wing discs were subjected to the immunofluorescence protocol described above. Following imaging, maximum projection images were processed with FIJI software (Schindelin et al., 2012). Using a box of identical dimensions, mean Arm signal intensity was quantified inside and outside of the transgene expression domain either on the most stabilized stripes of Arm protein or slightly above them as indicated. The ratio of signal inside/outside Arm signal was used to determine loss or gain or Arm levels within the transgene expression domain.

### Protein lysate preparation from larval heads and western blotting

Five late L3 developmentally stage-matched larval heads (with attached imaginal discs) were dissected and lysed with a 100 µl solution of 1× Cell Lysis Buffer (Cell Signaling Technology), supplemented with 1× Protease Inhibitors (Roche), 1 mM phenylmethylsulfonyl fluoride (PMSF) and 1 mM sodium fluoride (NaF). The tissues were mechanically homogenized and vortexed for 30 seconds. Lysates obtained after centrifugation for 10 min were mixed with 30 µl of 4X Laemmli buffer and the tubes were kept in boiling water for 5 minutes. The protein extracts were kept at –20°C until used for western blotting. Samples were then boiled for 5-10 minutes and then resolved on 10-12% SDS/PAGE gels before being transferred to nitrocellulose membranes. Following the transfer, membranes were blocked in 5% skimmed milk in TBS-T and then incubated with primary and secondary antibodies. The following primary antibodies were used: (1) mouse anti-FLAG (1:1000, Sigma M2), (2) mouse anti-β-tubulin (1:1000,104 Abcam G098). Anti-mouse HRP (1:5000, Jackson ImmunoResearch Laboratories) secondary antibody was used. Membranes were visualized using ClarityTM Western enhanced chemiluminescence (ECL) Substrate (BIO-RAD 170-5061), imaged on a GE AI 600 Imager and band density/protein levels were determined with FIJI software (Schindelin et al., 2012).

### Statistical analyses

Statistical analyses of the plotted data of mRNA and protein expression were done using one-way analysis of variance (ANOVA). For immunofluorescence experiments, paired (Arm stability within 1 wing disc) or unpaired (all other experiments) t-tests were performed to determine p-values. All statistical analyses were performed using GraphPad Prism 9.3.1 (GraphPad Software, San Diego, USA). Analyzed data with p<0.05 was considered statistically significant. Significance depicted as *p < 0.05, **p <0.01, ***p < 0.001, ****p < 0.0001, ns = not significant.

## Acknowledgements

We are grateful to the Developmental Studies Hybridoma Bank (IA, USA) for providing antibodies. We thank Dr. Janel Kopp for use of the 3DHistech slide scanner for all chicken histology images. We acknowledge the excellent virus injections carried out by Julian Kim and Adrian Danescu.

## Conflict of interests

The authors declare no competing or financial interests.

## Author contributions

Conceptualization: S.S.T and J.M.R.; Methodology: S.S.T and K.F.; Formal analysis: S.S.T. and J.M.R. Investigation: S.S.T., K.F., J.M.R.; Resources: J.M.R., E.M.V.; Writing - original draft: S.S.T. and J.M.R.; Writing - review & editing: S.S.T., J.M.R, E.M.V; Supervision: J.M.R.; Project administration: J.M.R., E.M.V.; Funding acquisition: E.M.V., J.M.R.

## Funding

This work was funded by the Canadian Institutes of Health Research (grant PJT-166182 to J.M.R. and E.M.V.). JMR holds a CRC Tier 1 chair (Award #CRC-2021-00441).

## Data availability

All relevant data can be found within the article and its supplementary information. The senior author JMR will provide additional data and all DNA constructs on request.

## Notes

### Competing Interest Statement

The authors have declared no competing interest.

