## supplementary data for "The abnormal C-terminus in DVL1 impacts Robinow Syndrome phenotypes"

### Supplementary figures

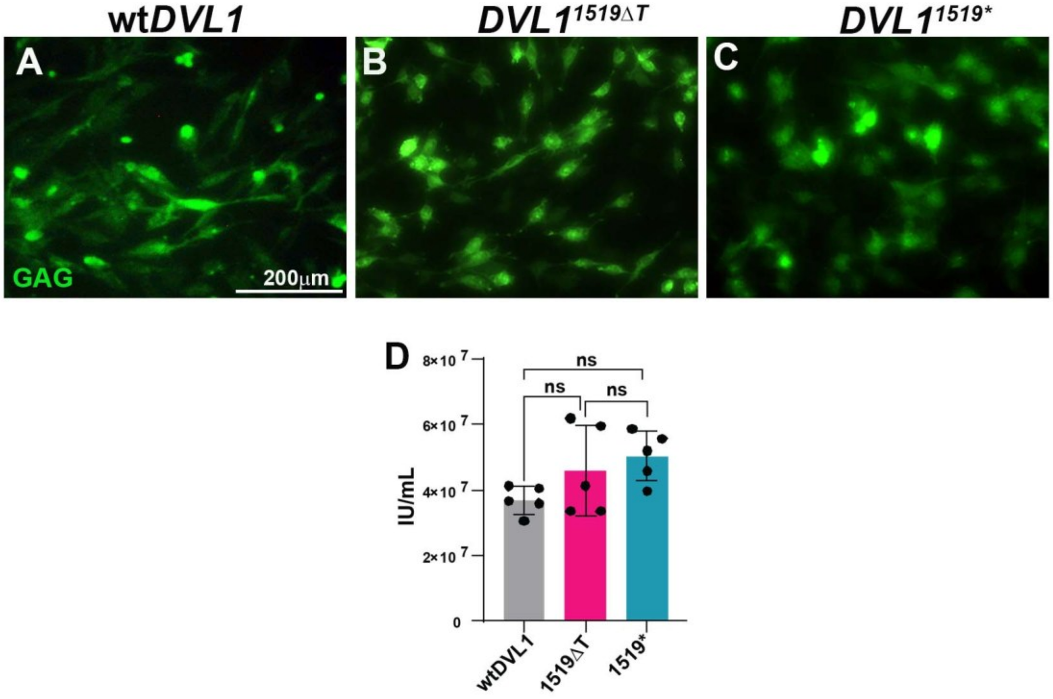


**Figure S1: Quantification of viral titer in cell cultures**

(A, A') Immunocytochemistry with anti-GAG (stains RCAS viral particles) performed on DF1 chicken fibroblasts infected with 2µl of high titre viral stock. Cells were fixed at 36h post-infection with virus. Counterstained with Hoechst. The GAG protein was detected with a mouse monoclonal antibody from Developmental Studies Hybridoma bank and detected with anti-mouse AlexaFluor 488. Quantification of infectious units (IU) showed that all h*DVL1* viruses had higher than the recommended titer of 1 x $10^7$IU/mL for performing overexpression studies. Scale bar = 200µm. Key: IU – Infectious units, ns – not significant.

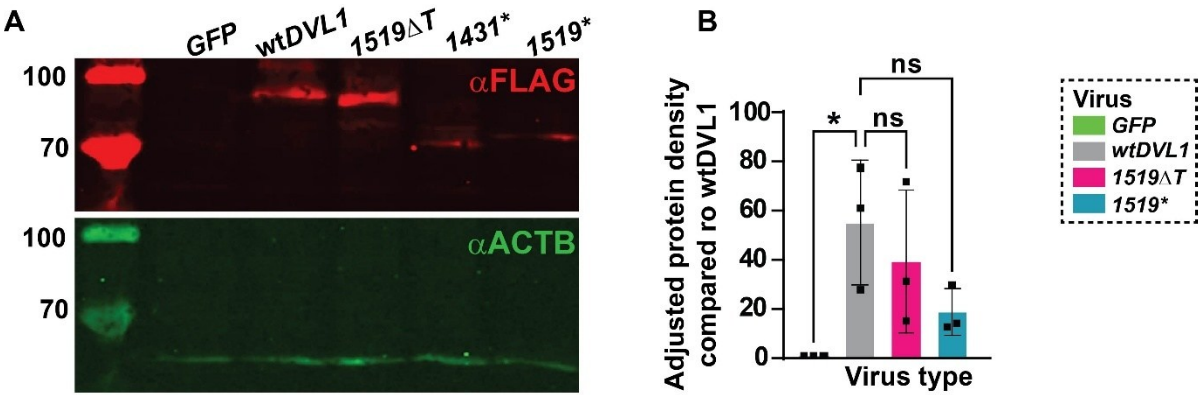


**Figure S2: Western blot analysis was conducted on DF1 chicken fibroblast lysate.**

The cells were transfected with FLAG-tagged hDVL1 construct and cultured for 4 weeks before lysis. (A) The Western blot probed with anti-FLAG antibody (Sigma, #F7425) shows comparable hDVL1 protein levels. Notably, there are expected size variations among the DVL1 proteins shown on the right side of the blot, wild-type DVL1 (~75 kDa), DVL1$^{1519\Delta T,\ 507fs*142}$ (~73 kDa), DVL1$^{1431*,\ 477*}$(~69 kDa), DVL1$^{1519*,\ 507*}$(~70kDa). β-actin (Abcam 1:2000 #8226, 42kDa) was used as a loading control. (B) Graph showing quantification of relative protein density compared to wtDVL1 (One-way ANOVA, Dunnett's test; n=3 independent blots). Statistical analysis was done in GraphPad Prism 10.1.0. Key: ns = non-significant.

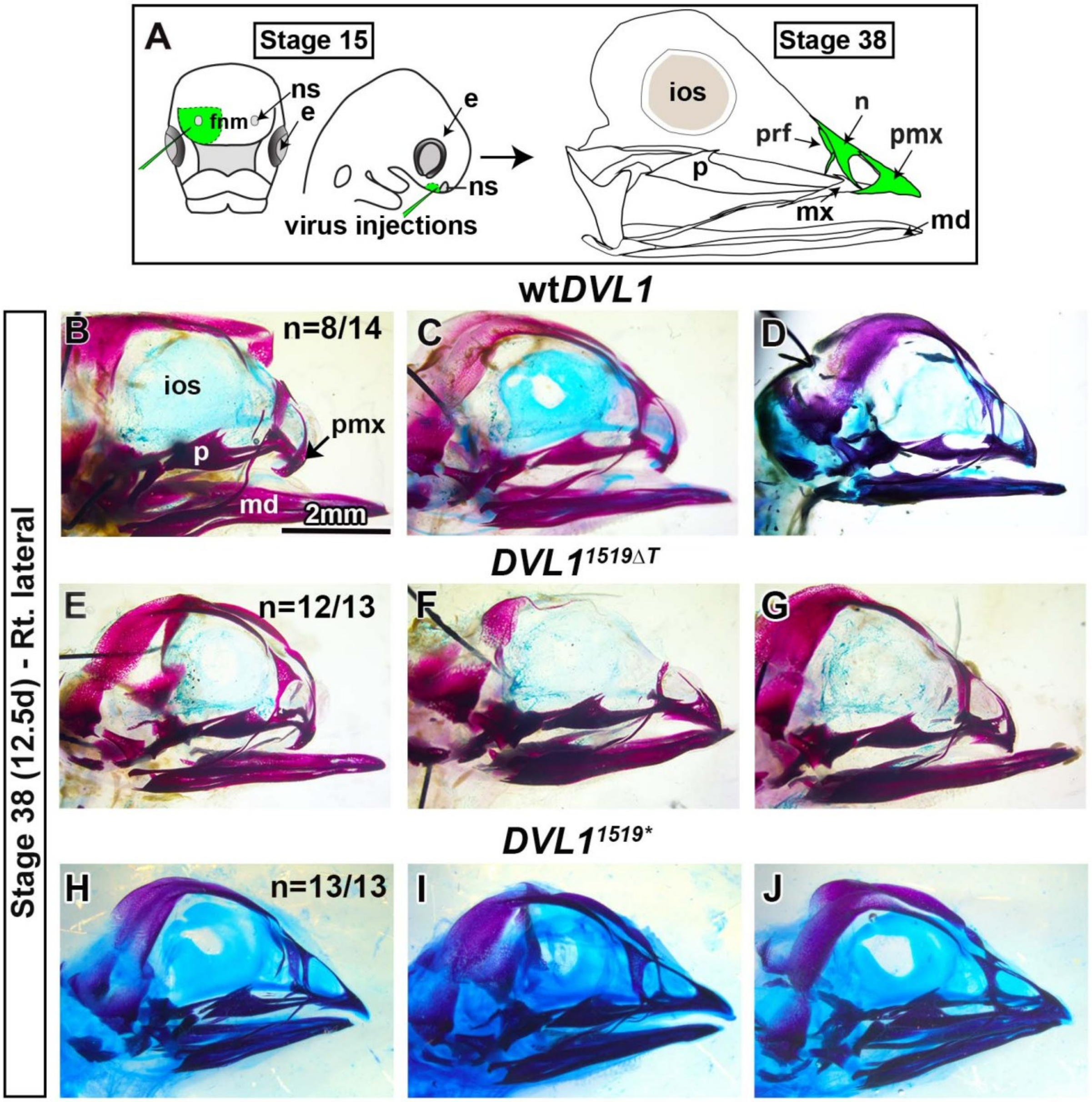


**Figure S3: Additional specimens treated with RCAS: wild-type h*DVL1* and RCAS:h*DVL1*$^{1519\Delta T}$**

Embryos injected with RCAS wild-type *DVL1* or *DVL1*$^{1519\Delta T}$ into the right frontonasal prominence at stage 15 (E2.5) and fixed 10 days post injection at stage 38 (E12.5). (A-C, D-F) External photographs in right lateral view of stage 38 embryos showing shortening and deviation of the upper beak caused by wild-type or mutant h*DVL1* viruses. (A'-C', D'-F') Wholemount skulls stained with alcian blue (for cartilage) and alizarin red (for bone) for the same specimens showing abnormal development of the skeletal elements. The faint cartilage stain (blue) in all skulls is a technical error. Scale bar: A-F' = 2mm.

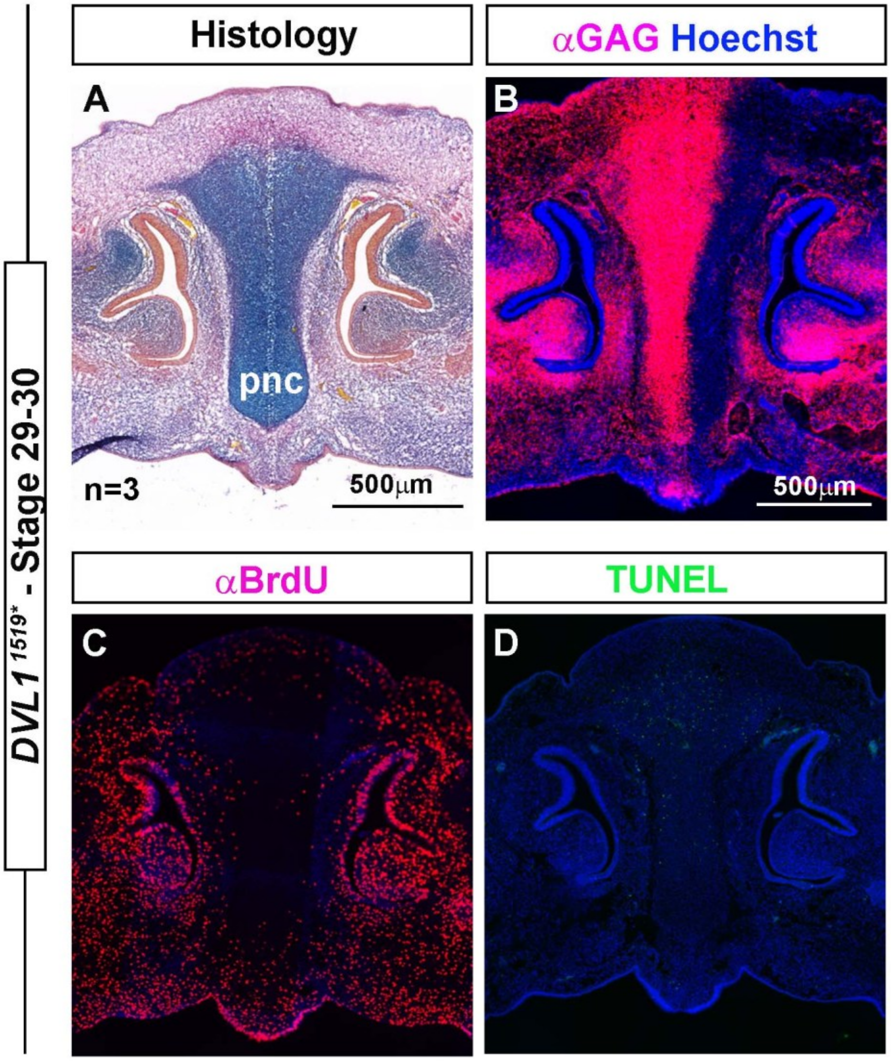


**Figure S4: Expression of the DVL1$^{1519*}$ did not affect proliferation or apoptosis**

Fixation of embryos approximately 96-100h post infection with virus and sectioned in the frontal plane shows A) normal histology of the prenasal cartilage. B) Robust GAG staining is present on the injected side of the frontonasal region. C) There were no qualitative effects on proliferation or on apoptosis (D). these data are consistent with the normal skeletal phenotype observed in Figure 1G-G''.

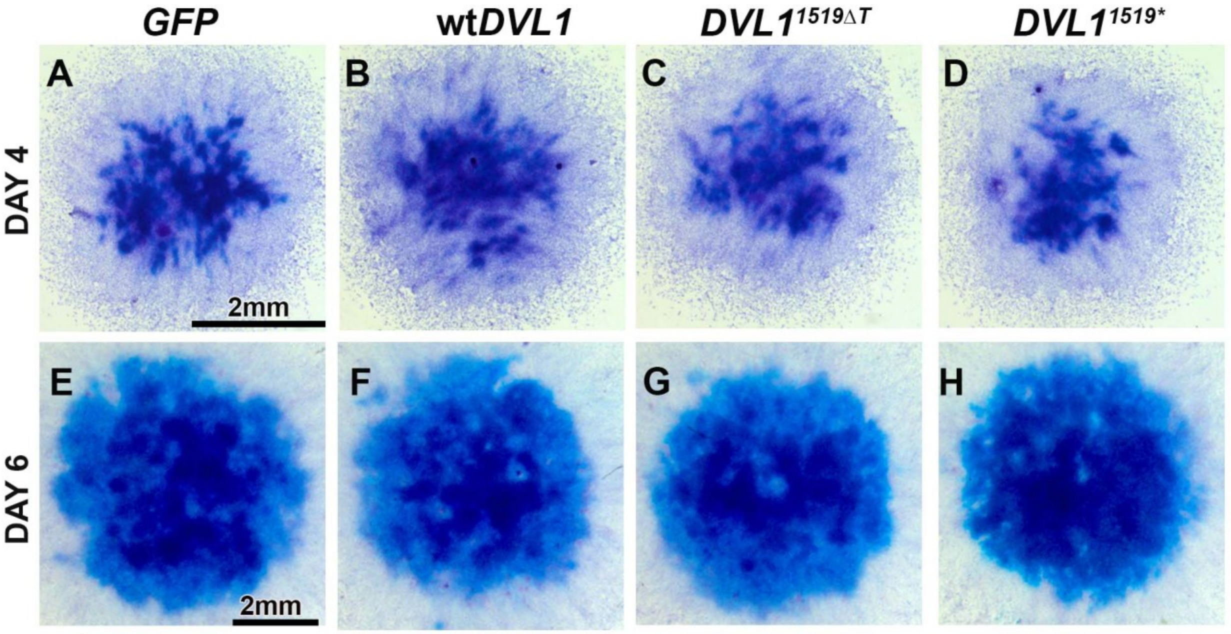


**Figure S5. Effects of h*DVL1* viruses on frontonasal mass micromass cultures**

Mesenchymal cells harvested from the frontonasal mass of stage 24 (E4.5) embryos were plated into high density cultures. Cultures were fixed and stained on day 4 (A-D), day 6 (E-H) or day 8 (I-L). The 8-day cultures were first stained with alkaline phosphatase to show mineralization (I-L) and then stained with Alcian blue for cartilage (I'-L'). Initiation of chondrogenesis was not affected. No measurable differences were detected between the different virus types after image analysis.

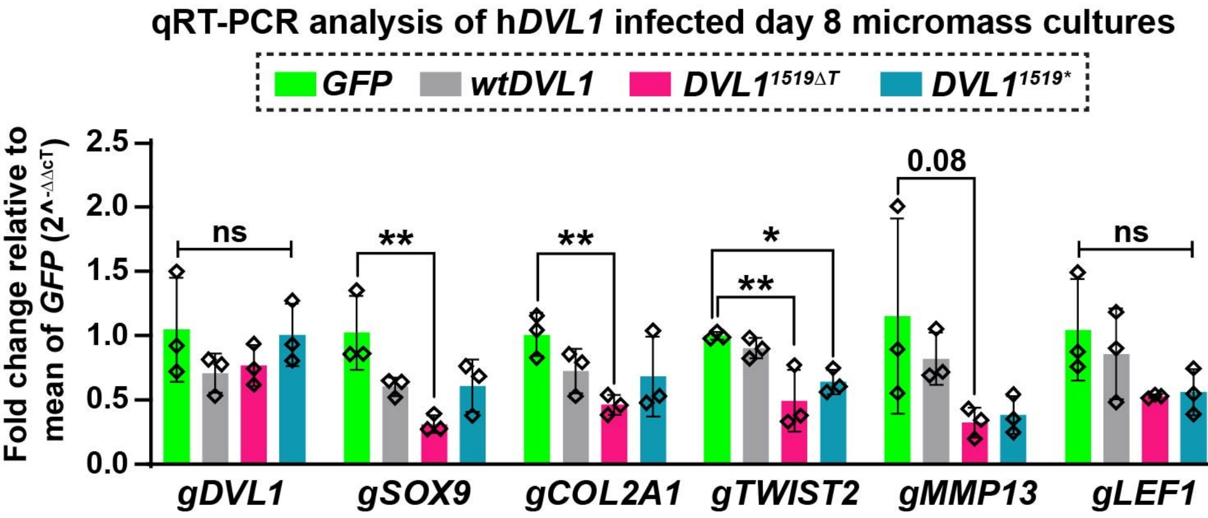


**Figure S6: Gene expression changes in 8-day micromass cultures infected with h*DVL1* viruses**

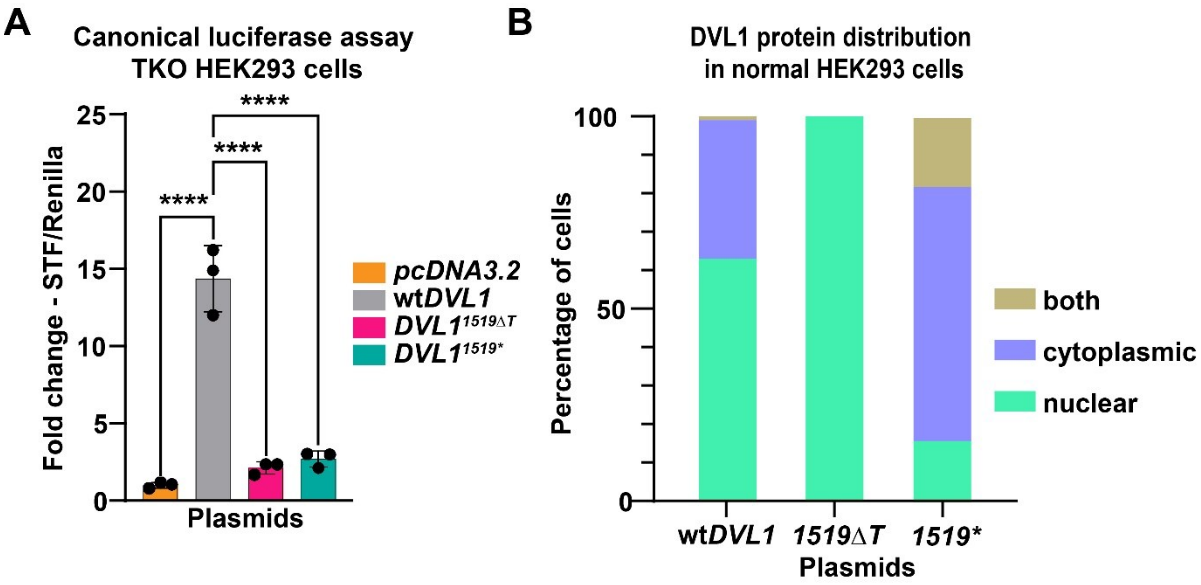


**Figure S7: Effects of DVL1$^{1519\Delta T}$ variant on WNT canonical pathway signaling and intracellular protein distribution**

A) The wtDVL1 plasmid strongly activates the SuperTopflash reporter however plasmids expressing the variant *DVL1$^{1519\Delta T}$* nor the *DVL1$^{1519*}$* truncated versions of the gene were able to activate the reporter. B) The same plasmids were transfected into wild-type HEK293T cells and stained with antibodies to the Flag tag. Distribution of protein expressed from the *DVL1$^{1519\Delta T}$* plasmid was restricted to the nucleus in contrast to the other forms of the plasmids.

#### Supplementary tables

**Table S1: Genes associated with the pathogenesis of Robinow syndrome**

| Gene/ OMIM ID | Human Chromosome locus | Inheritance | Function | Variant type | OMIM entry |
|---|---|---|---|---|---|
| ***WNT5A*** **164975** | 3p14.3 | Autosomal dominant | Ligand | Missense | 180700 |
| ***DVL1*** **601365** | 1p36.33 | Autosomal dominant | Cytoplasmic adaptor protein | -1 frameshift | 616331 |
| ***DVL2*** **602151** | 17p13.1 | Autosomal dominant | Cytoplasmic adaptor protein | +1 frameshift | No entry |
| ***DVL3*** **601368** | 3q27.1 | Autosomal dominant | Cytoplasmic adaptor protein | -1 frameshift | 616894 |
| ***FZD2*** **600667** | 17q21.31 | Autosomal dominant | Receptor-canonical and non-canonical WNT pathway | Missense, nonsense | No entry |
| ***ROR2*** **602337** | 9q22.31 | Autosomal recessive | Receptor – non-canonical WNT pathway | Missense, nonsense | 268310 |
| ***NXN*** **612895** | 17p13.3 | Autosomal recessive | Stabilizer of *DVL* | Deletion, missense | No entry |
| (Bunn et al., 2015; White et al., 2015; White et al., 2018; White et al., 2016; Zhang et al., 2022) | | | | | |

**Table S2 *DVL1* variants and patient phenotypes in autosomal dominant Robinow Syndrome**

| Total ADRS-*DVL1* patients | Male/ Female | Clinical phenotype | Type of variant (Frameshift) Reference sequence: NM_004421.3 |
|---|---|---|---|
| 23 | 11/12 | Frontal bossing 17/19, Midface hypoplasia 22/23 Hypertelorism 21/23 | c.1496_1508del, c.1505_1517del, c.1508del, c.1518del, c.1519del, c.1529del, c.1556del, c.1570_1571del, c.1608_1623del, c.1612_1615del, c.1615del, c.1623del, c.1619_1631del |

(Bunn et al., 2015; Hu et al., 2022; White et al., 2015; White et al., 2018; Zhang et al., 2022)

**Table S3 Sample size of embryos analyzed at each stage of development**

| Stage analyzed | Sample size (n) | Fixative | Experiment performed |
|---|---|---|---|
| **HH24 (E4.5, 48h)** | 3-6 | 4% PFA | Histology, immunostaining, qRT-PCR |
| **HH29 (E6.5)** | 3-6 | 4% PFA | Histology, immunostaining |
| **HH38 (E12.5)** | 13-35 | 100% ethanol | Skeletal staining & phenotypic analysis |

**Table S4: Prevalence of skeletal phenotypes produced by *DVL1* viruses**

| Virus type | Total embryos injected | % survival | Stage of collection | Total specimens (n) | Upper beak phenotype n, (%) | Premaxilla absent (n) |
|---|---|---|---|---|---|---|
| ***GFP*** | 35 | 69% | HH38 | 24 | 0 (0%) | 0 |
| **wt h*DVL1*** | 24 | 51% | HH38 | 14 | 8 (57%) | 8 |
| ***DVL1**$^{1519\Delta T}$ | 13 | 57% | HH38 | 13 | 12 (92%) | 12 |
| ***DVL1**$^{1519*}$ | 25 | 58% | HH38 | 13 | 0 (0%) | 0 |

**Table S5: Micromass cultures used for histology and immunostaining**

| Virus type | Total cultures collected for histology and immunostaining | |
|---|---|---|
| | **Day 6** | **Day 8** |
| ***GFP*** | 3 | 4 |
| **wt h*DVL1*** | 3 | 4 |
| ***DVL1**$^{1519\Delta T}$ | 3 | 6 |
| ***DVL1**$^{1519*}$ | - | 5 |

**Table S6: qRT-PCR primers**

| Gene | Accession # | Forward Primer | Reverse Primer |
|---|---|---|---|
| *hDVL1* | NM004421.3 | CAGCATAACCGACTCCACC | TGATGCCCAGAAAGTGATGTC |
| *gDVL1* | XM015297120.1 | CTCCCATTGAGAGGACAGGT | TGTTTCGTTGTCCAGTCCAT |
| *gWNT5A* | NM003392 | CAATGGCTTCTCAGTACCTCG | ACATCTGCACAGGGTTCATG |
| *gCTNNB1* | NM 205081 | CTTGGACTTGACATTGGTGC | CAGAGTGGAAAGAACGGTAGC |
| *gROR2* | NM 001080716.1 | AGTGCTGGAATGAATTTCCC | GGGAACTTGTTTGTGTGGTG |
| *gFZD2* | NM204222.2 | CTCCCATTGAGAGGACAGGT | TGTTTCGTTGTCCAGTCCAT |
| *gLRP5* | NM001012897 | ACCAAAGCCAGAACCCAG | CAGCACCATCCCTATTGACTC |
| *gATF2* | NM 204904 | GCCAGCGTTTTACCAATGAG | AGTTGGTGTTGGTGTCTGATC |
| *gLEF1* | NM_001398055.1 | CATCAAGTCCTCGCTGGTC | GCCCTTGTCATGGTAGGAATC |
| *gCOL2A1* | NM 204426.1 | GGACCAGCAAGACGAAAGAC | CGTAGCTGAAGTGGAAACCG |
| *gSOX9* | NM_204281.2 | CTGGGCAAGCTGTGGAG | GGTTGGTACTTGTAGTCGGG |
| *gTWIST1* | NM_204739 | GACTCCAAGATGGCAAGCTG | CTCCATTCTCCACACCGAGA |
| *gTWIST2* | NM_204679 | GAGTTATGCCTTCTCAGTCTGG | ACGTCCCAATTCCACTTCAG |
| *gBMP2* | NM204358 | GCTGTTTTGAGGTGGATTGC | AGGCACTGTTCTCTTTGTCC |
| *gBMP7* | XM 417496 | GTCAAACATCGCAGAGAACAG | TCCTTCACAGTAATACGCAGC |
| *gNOGGIN* | NM204123 | ACTTTATGGCTATGTCCCTGC | AACTCCAGCCCCTTGATTTC |
| *gBMPER* | NM 001007080 | AGCTGTCCTCATGGTAAAATCC | AACGTACTGACATGTTCCCTG |
| *gSP7* | XM015300329.3 | GTTCGTCTGCAATTGGCTCT | AATTTCTTCTCGCGGGTGTG |
| *gRUNX2* | NM204128 | ACCTAGTTTGTTCCCTGAACG | GTAATCTGACTCTGTCCTTGTGG |
| *gMSX1* | NM_204559 | TTCGGTCAAATCGGAGAACTC | TTCGTCTTGTGCTTCCTCAG |

**Table S7: Antibodies and immunofluorescence treatments**

<table>
<tr><th>Antigen Retrieval (steam 15 minutes, 95°C)</th><th>Permeabilization/ Pre-treatment</th><th>Blocking</th><th>Primary antibody, source, dilution (Overnight, 4°C)</th><th>Secondary Antibody and counterstain</th></tr>
<tr><td>10mM Sodium Citrate</td><td>0.1% Triton X-100 in 1XPBS for 15 min</td><td rowspan="8">10% Goat serum (Sigma G9023), 0.1% tween-20 in 1X PBS, (90 minutes, RT)</td><td>Ctnnb1/PY489-ß-catenin Developmental Studies Hybridoma bank (DSHB), 5µg/ml, Mouse monoclonal</td><td rowspan="8">ThermoFisher, Cy5 goat anti-mouse, #A10524,<br><br>ThermoFisher, Cy5 goat anti-rabbit, #A10523,<br><br>ThermoFisher, Alexa Fluor™ 488 goat anti-mouse, A11029,<br><br>ThermoFisher, Alexa Fluor™ 488 goat anti-rabbit A11034, 1:200 at RT in dark, 90mins<br><br>Hoechst (counterstain) 10µg/ml (Sigma 33568) in 1 X PBS (RT in dark for 30 min)</td></tr>
<tr><td>10mM sodium citrate</td><td>0.5% hyaluronidase in Hank's Balanced Salt Solution, 45min (following antigen retrieval)</td><td>Collagen type II DSHB, #II-II6B3, 5µg/ml, Mouse monoclonal</td></tr>
<tr><td>10mM Sodium Citrate</td><td>0.5% hyaluronidase in Hank's Balanced Salt Solution, 45min (following antigen retrieval)</td><td>Collagen type II Proteintech®, 28459-1-AP, 1:50, Rabbit polyclonal</td></tr>
<tr><td>1X Diva Decloaker (BioCare Medical)</td><td>0.1% Triton X-100 in 1XPBS for 15 min</td><td>Gag-pro AMV-3C2 DSHB, 1:4, Mouse monoclonal</td></tr>
<tr><td>10mM Sodium Citrate/ 1X Diva Decloaker (BioCare Medical)</td><td>0.1% Triton X-100 in 1XPBS for 15 min</td><td>SOX9 Sigma-Aldrich, #HPA001758 1:200, Rabbit Polyclonal</td></tr>
<tr><td>10mM Sodium Citrate/ 1X Diva Decloaker (BioCare Medical)</td><td>0.1% Triton X-100 in 1XPBS for 15 min</td><td>TWIST1 Twist2C1a Abcam #ab50887 1:50, Mouse monoclonal</td></tr>
<tr><td>0.3% Triton X-100 in 1XPBS for 15 min</td><td>0.1% Triton X-100 in 1XPBS for 15 min</td><td>Flag (DYKDDDDK) Thermo Fisher PA1-984B, 1µg/ml, Rabbit Polyclonal</td></tr>
<tr><td>1X Diva Decloaker (BioCare Medical)</td><td>0.1% Triton X-100 in 1XPBS for 15 min</td><td>BrdU DSHB, G3G4, 5µg/ml, Mouse monoclonal</td></tr>
</table>

**Table S8: List of primary antibodies used in the *Drosophila* study**

| Antibody | Dilution | Clone | Distributor |
| --- | --- | --- | --- |
| Mouse anti-FLAG | 1:500 | M2 | Sigma |
| Rabbit anti-FLAG | 1:500 | SIG1-25 | Sigma |
| Mouse anti-Armadillo | 1:50 | N2 7A1 | DSHB |
| Mouse anti-β-Galactosidase | 1:50 | 40-1a | DSHB |
